# Zika virus-specific IgM elicited during pregnancy exhibits ultrapotent neutralization

**DOI:** 10.1101/2021.11.23.469700

**Authors:** Tulika Singh, Kwan-Ki Hwang, Andrew S. Miller, Rebecca L. Jones, Cesar A. Lopez, Camila Giuberti, Morgan A. Gladden, Itzayana Miller, Helen S. Webster, Joshua A. Eudailey, Kan Luo, Tarra Von Holle, Robert J. Edwards, Sarah Valencia, Katherine E. Burgomaster, Summer Zhang, Jesse F. Mangold, Joshua J. Tu, Maria Dennis, S. Munir Alam, Lakshmanane Premkumar, Reynaldo Dietze, Theodore C. Pierson, Eng Eong Ooi, Helen M. Lazear, Richard J. Kuhn, Sallie R. Permar, Mattia Bonsignori

## Abstract

Congenital Zika virus (ZIKV) infection results in neurodevelopmental deficits in up to 14% of infants born to ZIKV-infected mothers. Neutralizing antibodies are a critical component of protective immunity. Here, we demonstrate that plasma IgM responses contribute to ZIKV immunity in pregnancy, mediating neutralization up to three months post symptoms. From a ZIKV-infected pregnant woman, we established a B cell line secreting a pentameric ZIKV-specific IgM (DH1017.IgM) that exhibited ultrapotent ZIKV neutralization dependent on the IgM isotype. DH1017.IgM targets a novel envelope dimer epitope within Domain II. The arrangement of this epitope on the virion is compatible with concurrent engagement of all ten antigen-binding sites of DH1017.IgM, a solution not achievable by IgG antibodies. DH1017.IgM protected against lethal ZIKV challenge in mice. Our findings identify a unique role of antibodies of the IgM isotype in protection against ZIKV and posit DH1017.IgM as a safe and effective candidate immunoprophylactic, particularly during pregnancy.

**Key points:**

- Plasma IgM contributes to early ZIKV neutralization during pregnancy
- Ultrapotent neutralization by pentameric DH1017.IgM mAb depends on isotype
- DH1017.IgM can engage all binding sites concurrently through different angles of approach
- DH1017.IgM protects mice against lethal ZIKV challenge

## Introduction

Zika virus (ZIKV) emergence in the Americas revealed it to be congenitally transmitted, causing microcephaly and other birth defects in up to 14% of infants born to ZIKV-infected pregnancies (Reynolds et al., 2017). Even infants born seemingly healthy following maternal ZIKV infection may develop neurodevelopmental defects years later (Nielsen-Saines et al., 2019). As ZIKV mostly results in mild febrile disease in healthy adults, its greatest disease burden arises through infections in pregnancy. In the 2015-2016 epidemic, ZIKV re-emergence in a susceptible population led to 11,000 cases of microcephaly in Brazil alone (Campos et al., 2018). There is no licensed vaccine for ZIKV, and the vaccine development and testing pipeline has stalled due to limited ZIKV circulation (Morabito and Graham, 2017).

Protective immunity in pregnancy could prevent ZIKV infection, congenital transmission, and lifelong congenital Zika syndrome sequelae. Neutralizing antibodies (nAbs) have been shown to be a critical aspect of protective immune responses against ZIKV and other flavivirus infections (Abbink et al., 2016; Hombach et al., 2005; Katzelnick et al., 2017; Kreil et al., 1997; Kwek et al., 2018; Larocca et al., 2016; Mason et al., 1973; Richner et al., 2017), yet nAb studies have largely focused on IgG antibodies. However, flavivirus infections are also characterized by virus-specific and prolonged IgM responses. IgM are pentameric molecules with 5-times as many potential antigen binding sites compared to IgG. Conventionally, IgM are thought of as early, short-lived, and low-affinity antibodies that are not somatically matured and thus unable to confer potent viral neutralization. Yet, early neutralizing IgM responses have been identified for other flaviviruses such as West Nile virus and yellow fever virus (Diamond et al., 2003; Gasser et al., 2020; Monath, 1971; Wec et al., 2020). A common feature is that these viruses present dense and repetitive structures on their surface, which may favor avid multivalent interactions with B cell receptors and influence the functional antibody profile via B cell stimulation and clonal selection. Interestingly, long-lasting ZIKV-reactive IgM responses have recently been independently identified in two distinct populations (Griffin et al., 2019; Stone et al., 2020). The presence of ZIKV-specific IgM lasting up to multiple years, when the typical half-life of IgM is 5 days, suggests that ZIKV-reactive IgM expressing B cells are specifically activated and expanded upon ZIKV infection (Lobo et al., 2004). While neutralizing activity is primarily attributed to IgG isotype antibodies, IgM may have an underappreciated role in ZIKV immunity.

The role of ZIKV IgM and IgM-producing B cells is especially understudied in pregnancy, a period of differential immunomodulation where ZIKV infections lead to their greatest disease burden. Early in pregnancy B cells are stimulated to produce IL-10, and subsequently B cell lymphopoiesis is suppressed, which promotes survival of mature B cells and decreases circulating naive B cell frequency (Christiansen et al., 1976; Lima et al., 2016; Nguyen et al., 2013). Retention of the IgM isotype diminishes as B cells differentiate from naïve to memory and antibody-secreting cells, which shapes B cell clonal selection in response to infection and long-lasting protective immunity (King et al., 2021). These factors may impact the magnitude and quality of ZIKV-specific IgM and IgG immunity in pregnancy.

In this study, we sought to define the kinetics and role of IgM in the control of ZIKV infection during pregnancy. We evaluated the contribution of plasma IgM to ZIKV neutralization in pregnant women over time and demonstrated that, in pregnancy, plasma IgM contributes to ZIKV neutralization primarily within the first 3 months of infection, regardless of prior exposure to other flaviviruses. We probed the B cell repertoire from peripheral blood of mothers with primary and secondary ZIKV infections and established 9 ZIKV-binding B lymphoblastoid cell lines (B-LCL). One of them produced an IgM antibody, DH1017.IgM, in its native pentameric conformation. DH1017.IgM was somatically mutated, did not cross-react with other flaviviruses, and displayed ultrapotent ZIKV neutralization that depended on its isotype. Structural studies identified a mode of antigen recognition on the ZIKV virion surface compatible with the concurrent engagement of all ten antigen-binding sites, a solution that is not available to antibodies of the IgG isotype, which have only two antigen-binding sites. The ultrapotent activity and protection mediated by DH1017.IgM in mice suggest that DH1017.IgM is a candidate for anti-ZIKV immunoprophylaxis.

## Results

### IgM contributes to plasma ZIKV neutralization during early phases of infection in pregnant women

We previously described a cohort of pregnant women from the 2015-2016 ZIKV outbreak in Brazil who presented with ZIKV-like symptoms of fever and/or rash in pregnancy and were serologically confirmed for ZIKV infection (Singh et al., 2019). Serologic evidence of prior exposure to DENV was demonstrated in most ZIKV-infected mothers in the cohort, and termed secondary ZIKV (Singh et al., 2019). Both primary (P24 and P54) and secondary (P14, P15, P17, P19, P23, P50, P56 and P73) ZIKV infection cases were included in this study. Among them, eight women were diagnosed during acute infection by serum PCR and two were identified as ZIKV-exposed based on ZIKV and DENV 1-4 neutralization titers (Singh et al., 2019). From 9 subjects, pregnancy and delivery plasma samples were available between 8- and 406-days post-symptoms (DPS). From 1 subject (P73), multiple longitudinal samples were collected between 8-406 DPS. Subject P73 developed prolonged viremia (up to 42 DPS), while all other subjects were viremic only up to their first visit (Singh et al., 2019). All subjects had high levels of ZIKV-neutralizing plasma antibodies at delivery (median FRNT_50_ = 4566; range: 650–14,959) (Singh et al., 2019).

To evaluate the contribution of plasma IgM to ZIKV neutralization during pregnancy, we measured total IgM and IgG concentrations, ZIKV binding IgG, and ZIKV neutralization of paired samples after mock and IgM depletion (**Table S1**). Subject P73 displayed the highest IgM-mediated ZIKV plasma neutralization, which started modestly at 8 DPS (7%), then rapidly peaked to 78% at 14 DPS ranging from 3% to 52% up to 112 DPS following a multimodal distribution and waned thereafter to undetectable levels. (**Figure 1**). An important contribution of IgM to plasma ZIKV neutralization was confirmed across all the samples collected between 14 and 100 DPS from each of the other subjects (range: 17-49%), regardless of serostatus (i.e., primary or secondary ZIKV infection). IgM plasma contribution was minimal to null in samples collected during late infection (range: 0-4%), except for subject P23, who displayed 28% of IgM-mediated ZIKV neutralization at 209 DPS. Overall, 90% (9/10) of plasma samples collected within 3 months of symptomatic infection from 4 mothers demonstrated >10% IgM-attributable ZIKV neutralization, but only 18% (2/11) of samples from beyond 3 months demonstrated this response (**Figure 1**).

**Figure 1.**
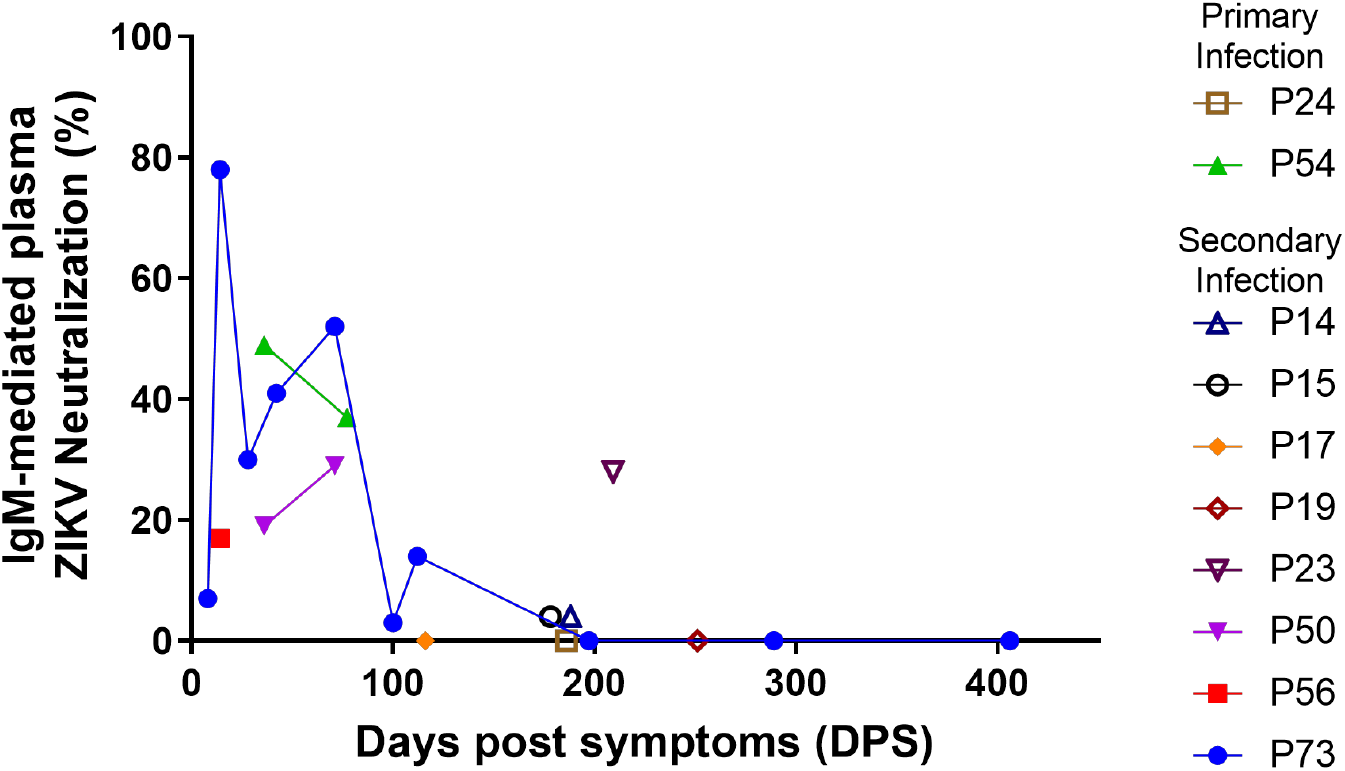
Contribution of plasma IgM to ZIKV neutralization from acute infection through late convalescence. Percent ZIKV neutralization attributable to plasma IgM across ZIKV-infected mothers (n=10) from 8 to 406 days post symptoms (DPS). Each sample tested in three technical replicates. See also **Table S1**.

### Isolation of ZIKV-specific, ultrapotent neutralizing mAb DH1017.IgM

We next probed the B cell compartment of subject P73 at 28 and 71 DPS to identify B cells producing ZIKV-neutralizing IgM. We also selected four ZIKV-infected pregnant women from the same cohort (Singh et al., 2019), including one woman with primary ZIKV infection (P54: 29 and 77 DPS) and three women with secondary ZIKV infection (P34 at 162 DPS, P56 at 19 DPS, and P73 at 28 and 71 DPS) for a comparative analysis of the ZIKV-specific B cell repertoire. We enriched for ZIKV-reactive B cells by sorting either unfractionated or memory B cells (MBCs) using a fluorescent ZIKV virion as bait and cultured cells at limiting dilution to derive human B-LCL through EBV transformation as described (Bonsignori et al., 2011, 2018).

Across subjects, ZIKV-reactive B cells constituted 0.6-1.7% of the circulating unfractionated B cell pool (**Figure S1A**). Regardless of serostatus and time of sample collection relative to symptoms onset, the frequency of ZIKV-reactive MBCs out of total MBCs was uniformly higher than the ZIKV-reactive B cells out of the total unfractionated B cell pool (0.7-2.6% vs 0.6-1.7%; p<0.03, paired t-test) (**Figure S1A**). On average, ZIKV-reactive memory B cells comprised 52% of total ZIKV-reactive B cells with higher frequencies observed in secondary infections (P73, P56, P34) than in the primary ZIKV infection case (P54) (46-69% vs 34%; **Figure S1B**), compatible with a re-engagement of pre-existing cross-reactive memory B cells in secondary infection.

In vitro B cell stimulation across all 4 subjects yielded a total of 97 Ig-secreting culture supernatants, of which 37 were from unfractionated B cells and 60 from memory B cells. Among unfractionated cultures, 13 cultures contained both IgG and IgM, whereas all cultures from memory B cells were of a single Ig isotype. Ig concentrations ranged from 3 to >3000 ng/ml with a geometric mean of 146 ng/ml. Overall, 49 of 97 (50.5%) Ig-secreting culture supernatants confirmed ZIKV reactivity: 17 (15 IgG and 2 IgM) from unfractionated B cells and 32 (all IgG) from memory B cell cultures (**Figure S1C**). From ZIKV-reactive Ig-secreting cultures, we established 9 B-LCL, 8 of which produced IgG monoclonal antibodies (mAbs) and one, termed DH1017.IgM, produced an IgM mAb.

Analysis of the Ig V(D)J rearrangement immunogenetics demonstrated that none of the B-LCLs were clonally related (**Table 1**). Diverse heavy chain variable gene segments were used and paired with either κ or λ light chains (V_L_), without V_H_/V_L_ pairing preferences. CDRH3 lengths ranged between 12 and 22 amino acids (median: 15 amino acids). Overall, V_H_ and V_L_ somatic hypermutation (SHM) frequency ranged from 3.1% to 10.8%, and 1.5% to 8.0%, respectively (**Table 1**). The V_H_ SHM frequency of sequences obtained from unfractionated B cells and memory B cells was statistically similar (p>0.05, Kolmogorov-Smirnov test). DH1017.IgM used the V_H_4-31 gene segment paired with V_λ_ 1-51 and had a 15 amino acid-long CDRH3 (**Table 1**). While the B cell that expressed DH1017.IgM was isolated from unfractionated B cells and we could not determine its origin from the naïve or memory B cell compartment, the SHM frequency in the DH1017.IgM V(D)J rearrangements were within the range of other IgG mAbs that we isolated (3.8% for the heavy chain and 3.4% for the light chain) with 7 nucleotide mutations in the V_H_ and 9 in the V_L_. All but two of these nucleotide changes were substitution mutations. Nine of the 16 nucleotide substitutions (56%) occurred in canonical activation-induced cytidine deaminase (AID) hotspot motifs with high mutability rates (Yaari et al., 2013). The remaining 7 substitutions occurred in neutral or cold-spot motifs (**Table S2**).

**Table 1.**
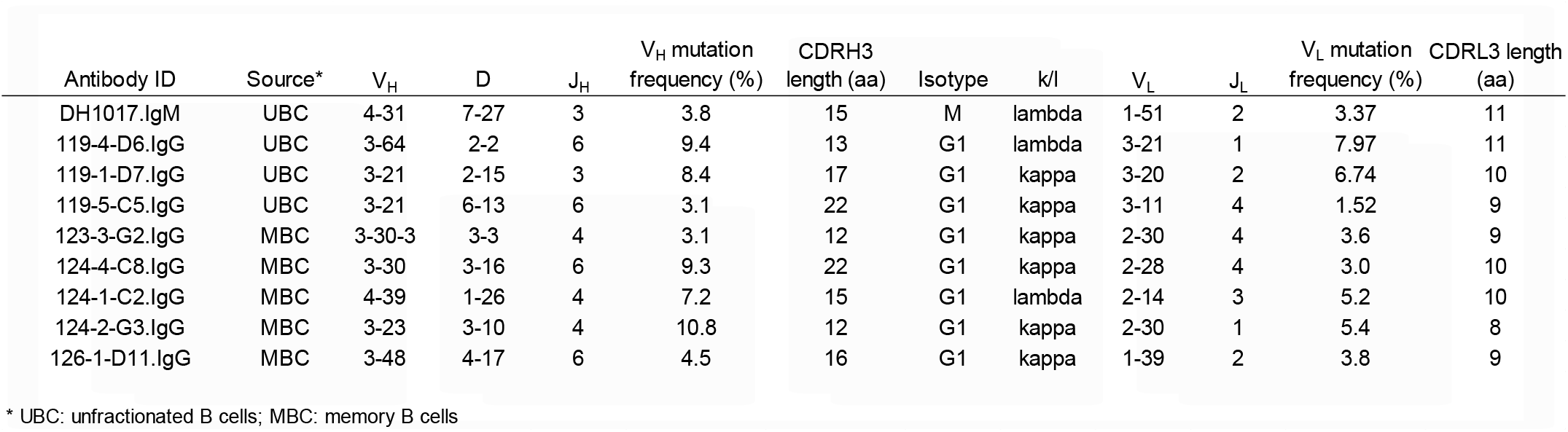
Immunogenetics of monoclonal antibodies isolated from B-LCLs. See also **Figure S1** and **Table S2**.

MAbs were produced and purified from the 9 B-LCLs for functional characterization. In a native gel, mAb DH1017.IgM yielded two bands at ∼970 kDa and ∼1048 KDa (**Figure 2A**), which is compatible with pentameric and hexameric IgM isoforms (Keyt et al., 2020; Wiersma et al., 1998). Negative stain electron microscopy class average analysis supported the presence of both pentameric and hexameric isoforms, with hexamers representing 18% and pentamers representing 73% of all observed images (**Figure 2A**).

**Figure 2.**
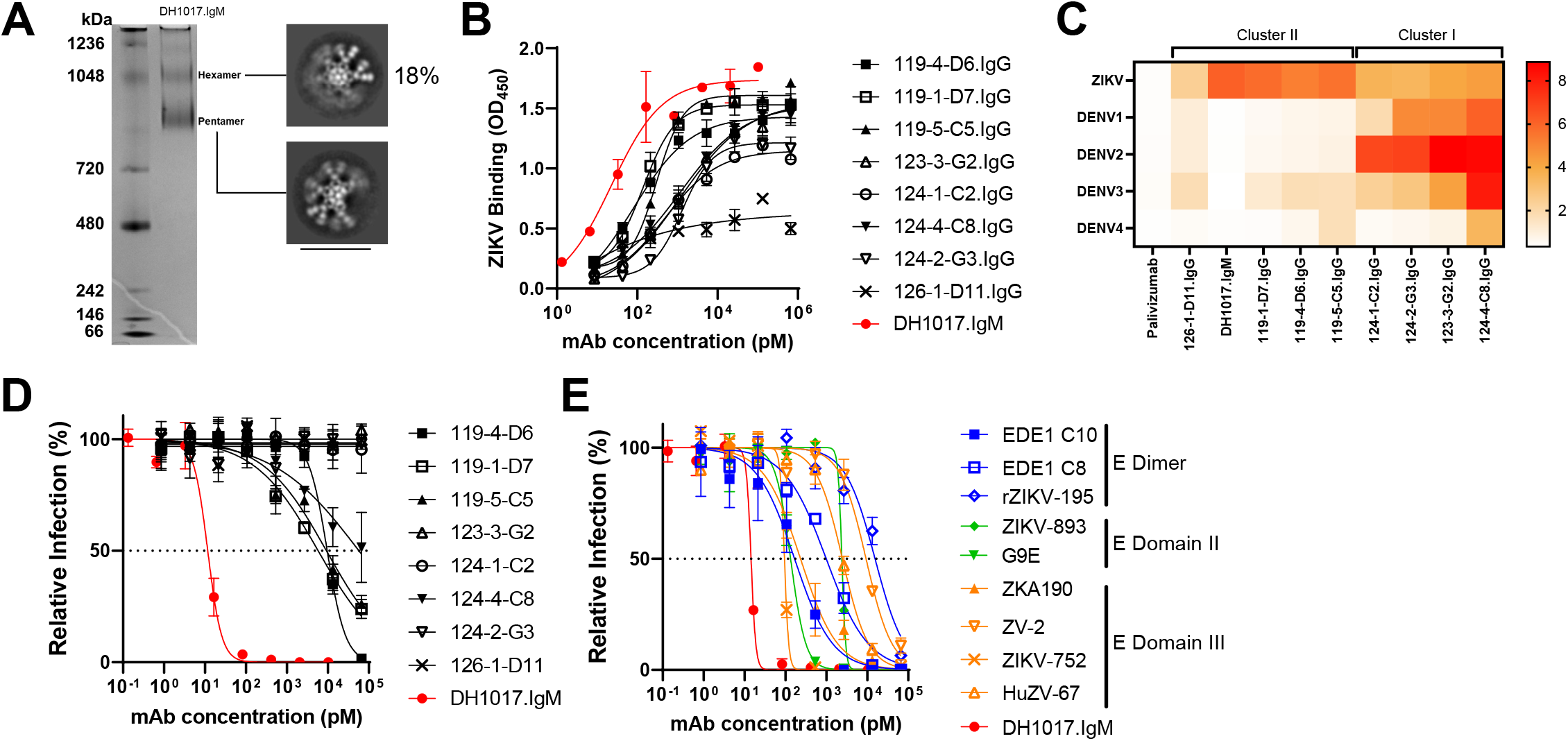
Characterization of B-LCL-derived ZIKV-specific monoclonal antibodies. **A.** Left: DH1017.IgM native PAGE gel in non-reducing conditions. Bands identified high and low sized molecular species compatible with IgM hexameric and pentameric forms. Ladder on the left. Right: Negative stain electron microscopy of purified DH1017.IgM showing representative class averages of hexameric and pentameric particles. Scale bar is 40nm. **B.** Binding to whole ZIKV PRVABC59 virions by B-LCL-derived mAbs (n=9). DH1017.IgM is shown in red. Error bars indicate standard deviation of two technical replicates. Experiments were independently repeated twice. **C.** Heatmap showing binding, expressed as LogAUC, to ZIKV and DENV1-4 by the nine B-LCL-derived mAbs. Palivizumab is shown as negative control. Clustering was performed using the Los Alamos Database heatmap tool. **D.** ZIKV PRVABC59 strain neutralization curves of the 9 B-LCL-derived mAbs expressed as percentage of the number of foci relative to the virus alone condition. Dotted line represents 50% viral inhibition. DH1017.IgM is shown in the red filled circle. Error bars indicate standard deviation of assays run in triplicate. Experiments were independently repeated twice. **E.** ZIKV PRVABC59 strain neutralization curves of DH1017.IgM (red) and 9 previously described IgG ZIKV neutralizing antibodies. Colors indicate groups of similar epitopes. Error bars show standard deviation in triplicate measurements. Experiments were independently repeated twice. See also **Figure S2**.

All purified mAbs confirmed binding to ZIKV. While 126-1-D11.IgG bound weakly even at high concentrations (LogAUC = 2.0), all other mAbs bound with LogAUC ranging from 2.7 to 5.9, with the strongest ZIKV-binding mAb being DH1017.IgM (**Figure 2B**). Since plasma antibody displayed substantial cross-reactivity with DENV in all subjects except primary infection subject P54 (**Figure S2A**), we determined cross-reactivity of the nine mAbs with DENV 1-4 strains. Based on their binding profile, the mAbs segregated into two clusters (**Figure 2C**). Cluster I comprised mAbs 123-3-G2.IgG, 124-4-C8.IgG, 124-1-C2.IgG, and 124-2-G3.IgG that bound to one or more DENV 1-4 strains better than to ZIKV. These mAbs were isolated from memory B cells of mothers with secondary ZIKV-infection (P73 and P56), which parallels the plasma cross-reactivity profile and further implies re-engagement of pre-existing DENV-reactive memory B cells. Conversely, cluster II mAbs bound more strongly to ZIKV than to DENV, for which cross-reactivity was either at or below limit of detection, which suggests that these mAbs may constitute a *de novo* immune response to ZIKV. Notably, DH1017.IgM did not cross-react with or neutralize any of the four DENV serotypes (**Figure 2C** and **Figure S2B**).

Five of the nine ZIKV-specific mAbs mediated neutralization of the ZIKV PRVABC59 strain (**Figure 2D**). Mab 124-4-C8.IgG neutralized weakly with FRNT_50_ >10µM. Mabs 119-1-D7.IgG, 119-5-C5.IgG and 119-4-D6.IgG neutralized with FRNT_50_= 5.8 nM, 9.2 nM and 9.7 nM, respectively. Remarkably, DH1017.IgM neutralized ZIKV with ∼500- to 1000-fold higher potency (FRNT_50_=12 pM) (**Figure 2D**). DH1017.IgM neutralization of the ZIKV PRVABC59 strain was repeated multiple times (n=10) and potent neutralization was independently confirmed by multiple operators (geometric mean [GM] FRNT_50_: 12 pM; range: 4-31 pM). DH1017.IgM also neutralized the ZIKV H/PF/2013 strain with comparable potency (GM FRNT_50_: 14 pM_;_ range: 8-26 pM) (**Figure S2C**). We compared DH1017.IgM potency with a panel of well characterized ZIKV-neutralizing IgG mAbs: EDE1 C8, EDE1 C10, ZIKV-893, rZIKV-195, G9E, ZV-2, ZIKV-752, ZKA190, and HuZV-67 (Collins et al., 2019; Gilchuk et al., 2020; Hasan et al., 2017a; Long et al., 2019; Rouvinski et al., 2015; Sapparapu et al., 2016; Wang et al., 2017; Zhao et al., 2016). In this side-by-side comparison, DH1017.IgM neutralized more potently than all other IgG ZIKV neutralizing mAbs in the panel (FRNT_50_ range: 95 pM – 15.5 nM) (**Figure 2E**). Neutralizing antibodies of the IgG isotype with IC_50_ < 10ng/ml, i.e. 66.7pM, are defined as ultrapotent (Smith et al., 2015). Hence, DH1017.IgM meets the definition of ultrapotent ZIKV-neutralizing antibody.

### Impact of isotype on DH1017.IgM functions

We recombinantly produced the Fab and IgG1 isoforms (DH1017.Fab and DH1017.IgG) and measured binding to ZIKV H/PF/2013 envelope (E) protein monomer and dimer. DH1017.IgM bound more potently than DH1017.IgG and DH1017.Fab to ZIKV E monomer (EC_50_ = 202 pM, 463 pM and 1.24 nM, respectively) (**Figure 3A**). The difference in binding potency among the different isotypes was more profound for the ZIKV E dimer: DH1017.IgM EC_50_ = 91 pM, DH1017.IgG EC_50_ = 929 pM, and DH1017.Fab EC_50_ = 25.1 nM (**Figure 3B**). To test the contribution of the Fc region of DH1017.IgM to its increased level of binding, we compared the DH1017.IgM and DH1017.IgG binding kinetics to ZIKV E dimer using a BLI format with immobilized mAb. Under these conditions, DH1017.IgM and DH1017.IgG displayed similar dissociation kinetic (k_off_ = 1.03 and 0.91 s^-1^x10^-2^, respectively) and affinity (K_d_ = 106 and 123 nM, respectively). This suggests that the µ-chain Fc of DH1017.IgM does not contribute towards its affinity for E protein.

**Figure 3.**
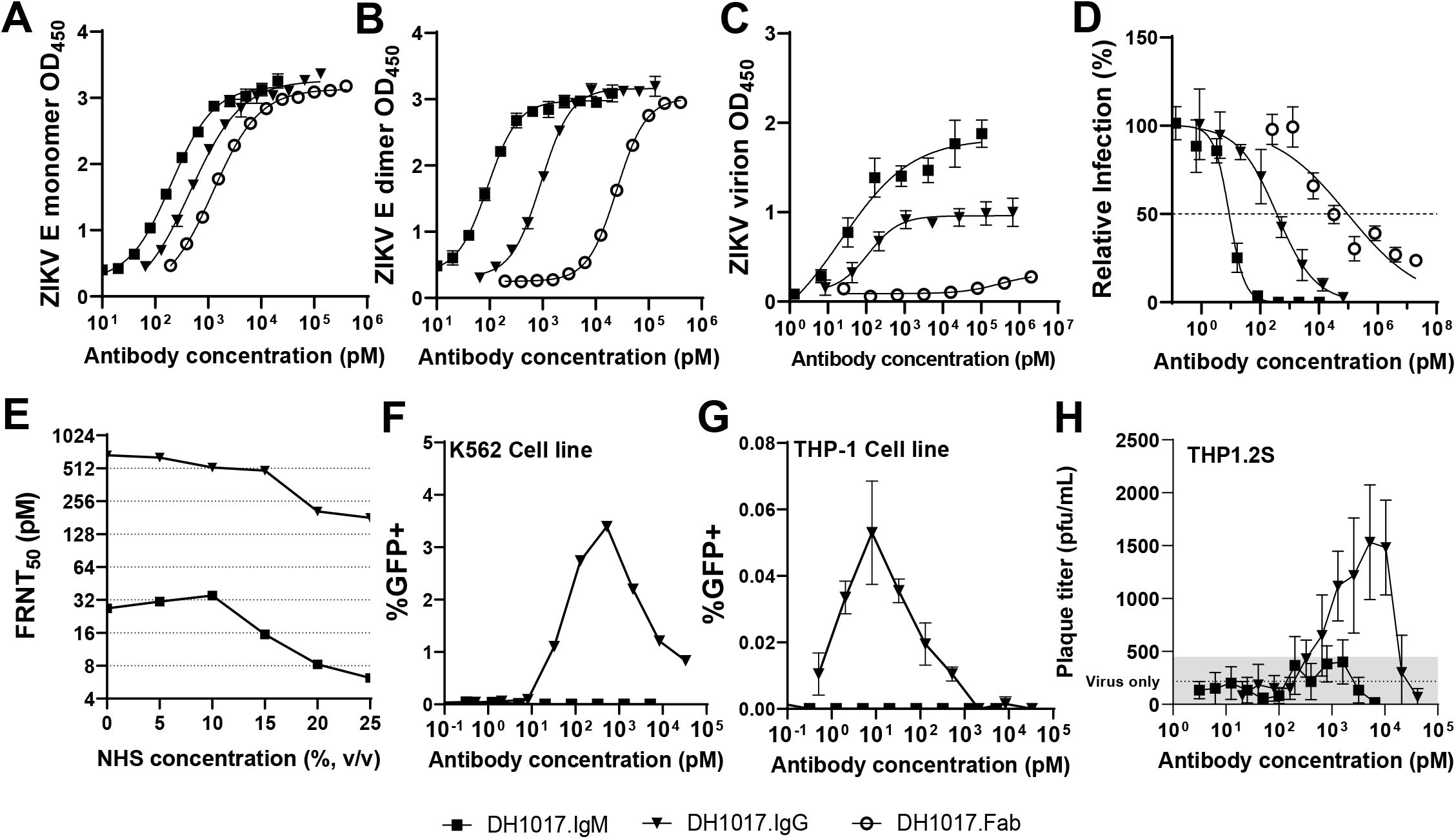
Functional characterization of DH1017.IgM, DH1017.IgG, and DH1017.Fab. **A, B.** Binding of DH1017.IgM (square), DH1017.IgG (triangle) and DH1017.Fab (circle) mAbs to ZIKV E monomer (A) and dimer (B) measured by ELISA and expressed as optical density at 450nm (OD_450_). Error bars indicate standard deviation of duplicate observations. Experiments were independently repeated twice. **C.** Whole ZIKV virion binding curves of DH1017.IgM, DH1017.IgG, and DH1017.Fab. Error bars show the standard deviation of two experiments, each run with 6-replicates for DH1017.IgM and DH1017.IgG. DH1017.Fab was run in duplicate. **D.** ZIKV neutralization of serially diluted mAbs. Dotted line denotes 50% relative infection of mAb condition as compared to virus alone (FRNT_50_). Error bars indicate standard deviation of mAbs run in triplicate. Experiments were independently repeated twice. **E.** ZIKV neutralizing titers at 50% viral inhibition (FRNT_50_) over increasing complement concentrations. Normal human serum (NHS) was used as source of complement at the indicated concentrations (v/v). Percent infection of the antibody were calculated relative to the virus alone with matched concentrations of NHS. MAbs were run in triplicate. **F, G**. Serially diluted mAbs were incubated with ZIKV H/PF/2013 reporter virus particles and used to infect K562 (**F**) or THP-1 (**G**) cells to measure antibody-dependent enhancement (ADE) of infection. Enhancement of infection was scored by GFP expression and measured after 36-48 hours using flow cytometry. Representative ADE data of at least three independent experiments are shown. **H.** ADE of DH1017.IgM and DH1017.IgG on THP1.2S cells measured by plaque assay. Dotted line shows mean of six virus-only control replicates and the grey area indicates one standard deviation above and below this mean. Representative results from duplicated experiments run with 6-replicates each. Bars show the standard deviation from 6 replicates. See also **Figure S3**.

We then assessed the ability of DH1017.IgM, DH1017.IgG and DH1017.Fab to bind to whole Zika PRVABC59 virion. DH1017.Fab only interacted weakly with ZIKV (EC_50_ >2mM) whereas DH1017.IgG bound strongly (EC_50_ = 112 pM), and DH1017.IgM bound 5-fold more potently than DH1017.IgG (EC_50_ = 22 pM) (**Figure 3C**). Remarkably, the functional difference between DH1017.IgM, DH1017.IgG and DH1017.Fab was even greater for ZIKV neutralization. DH1017.IgM neutralization potency (FRNT_50_ = 9 pM) was >40-fold higher than DH1017.IgG (FRNT_50_ = 366 pM; **Figure 3D**), whereas DH1017.Fab yielded a shallow neutralization curve (FRNT_50_ = 93 nM) with 10,000-fold worse potency than DH1017.IgM (**Figure 3D**). These data demonstrate that DH1017.IgM increased binding and ultrapotent neutralization depended on the IgM isotype.

Since antibody-mediated complement deposition can reduce the amount of mAb required to neutralize virus, we tested the effect of complement from normal human serum (NHS) on DH1017.IgG and DH1017.IgM neutralizing activities. Exogenous human complement from NHS enhanced ZIKV neutralization potency of both mAbs in a dose-dependent manner (**Figure 3E**). At all doses of NHS tested (5-25% v/v), DH1017.IgM retained more potent neutralizing activity compared to DH1017.IgG. The largest improvement in neutralization potency was observed in the 25% NHS condition with 4.3-fold and 3.7-fold increase in potency for DH1017.IgM and DH1017.IgG, respectively (**Figure 3E**). These data demonstrate that neutralization potency of the DH1017 mAb in its IgG and IgM isoforms is enhanced in presence of complement.

Sub-neutralizing concentrations of flavivirus neutralizing IgG can mediate antibody-dependent enhancement (ADE) of viral infection through interactions of immune complexes with plasma membrane-anchored FcγR, particularly on monocytes (Chan et al., 2015; Halstead, 1988; Halstead and O’Rourke, 1977). DH1017.IgG mediated ADE in both K562 and THP-1 cells whereas DH1017.IgM did not (**Figures 3F, G**). DH1017.IgM also did not mediate ADE in THP-1.2S cells, a subclone of THP-1 cells with increased sensitivity to ADE (Chan et al., 2014) (**Figure 3H**). Similarly, DH1017.IgM did not mediate ADE in primary monocytes (**Figure S3A**). Finally, to further characterize the safety profile of DH1017.IgM, DH1017.IgG and DH1017.Fab, we measured binding to nine autoantigens associated with autoimmune diseases. All isotypes tested negative for all antigens (**Figure S3B**). Further, they did not demonstrate intracellular immunofluorescent staining of Hep-2 cells (**Figure S3C**).

### Structural characterization of the epitope of the DH1017 clone on the Zika virion

Using cryo-electron microscopy (cryo-EM) and single particle reconstruction, we identified a set of 4104 ZIKV H/PF/2013 particles bound with DH1017.Fab and generated a cryo-EM density map. A surface shaded view of the virus-Fab complex is shown in **Figure 4A** and demonstrates the sites of Fab binding to the surface of the particle. The cryo-EM density map resolution is 5.3 Å as determined from the Fourier Shell Correlation co-efficient 0.143 (**Figure S4A**).The C_α_ backbone of the E glycoprotein ectodomains from the ZIKV asymmetric unit (PDB 6CO8; Sevvana et al., 2018) is composed of three E monomers, chains A, C, and E (**Figure 4B**). Their position in the density map is shown in **Figure S4B**. In this asymmetric unit, Chain A lies alongside the antiparallel homodimer formed by chains C and E (**Figure 4B**). The fitted asymmetric unit is repeated 60 times within the density map demonstrating icosahedral symmetry like previous Zika virus structures (Sevvana et al., 2018; Sirohi et al., 2016).

**Figure 4.**
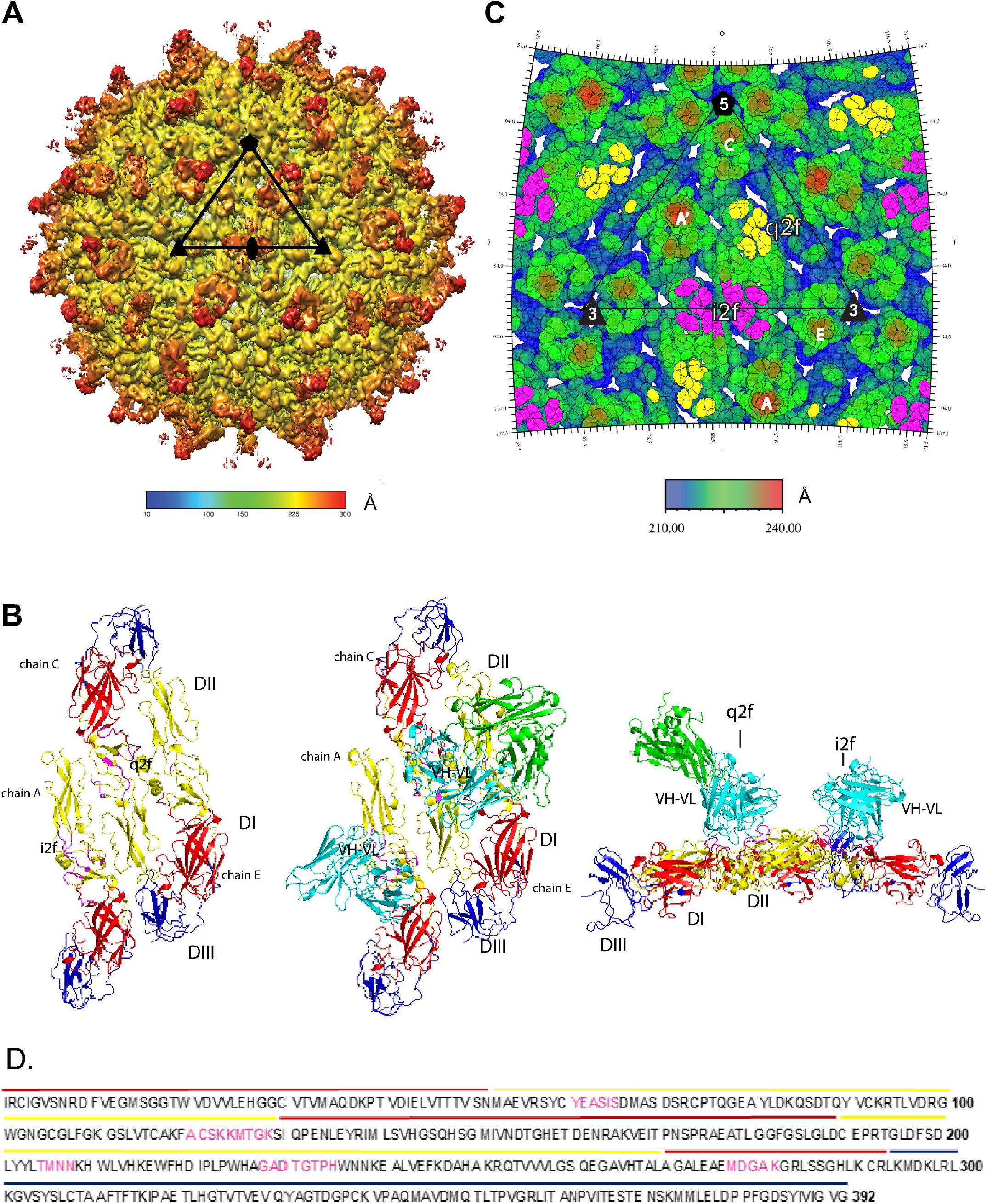
DH1017 clone interacts with envelope dimer. **A.** Surface-shaded view of the Zika virion bound with the DH1017 Fab fragment at a resolution of 5.3 Å acquired via cryo-EM. The color represents the distance to the center and shown by the colored scale bar in Å. The black triangle represents the asymmetric unit, a pentagon represents the five-fold axis, the triangle the three-fold axis, and the oval the two-fold axis. **B.** Surface representation of the E ectodomain of Zika virus asymmetric unit (PDB 6CO8) shown in top view unbound, then bound with the variable domain of DH1017.Fab (cyan) and the constant domain (green), and a side view of the asymmetric ectodomain bound the DH1017 Fab fragment (left to right). The E ectodomains are colored with DI in red, DII in yellow, and DIII in blue. The residues of the footprint are colored, magenta. See also **Fig. S4 and Table S3**. **C**. A radially colored roadmap (scale bar in Å). The icosahedral and quasi two-fold axes are labelled i2f and q2f, respectively. The monomer chains of two E dimers are labeled A’ and A at the i2f, and chains C and E at the q2f. Residues on the surface of the virus within 6 Å of the variable domain structure fit to the density map are colored yellow and magenta. Yellow residues are on chain C at the q2f axis and magenta residues are at the i2f axis on chain A’ and A. **D**. Fab DH1017 epitope shown on the primary sequence of E ectodomain. The epitope residues are colored magenta and the domains DI, DII, and DIII are indicated by line color red, yellow and blue, respectively.

The C_α_ backbone of the homology model of the DH1017.Fab was fit into the cryo-EM density map (**Figure S4B**). It shows that the DH1017 variable domain interacts primarily with DII of all three E monomers in the asymmetric unit, and at the interface of DII and DI on chains A and C (**Figure 4B, S4B**). The DH1017.Fab epitope footprint was defined by residues of the E ectodomain glycoprotein on the surface of the virus within 6 Å from the C_α_ backbone of the DH1017 variable domain structure (**Table S3, Figure 4B, C, D**). The DH1017.Fab paratope footprint was defined by Fab residues within 6 Å from the C_α_ backbone of the E ectodomain glycoprotein of the virus surface and included all three heavy chain CDRs and FRH3 (**Table S3**).

There are two 2-fold axes of symmetry on the virus surface: the icosahedral 2-fold (i2f) axis delineates symmetry at the juncture of two antiparallel asymmetric units on the repetitive virion surface between A and A’, whereas the quasi 2-fold (q2f) axis describes symmetry within the antiparallel E dimer between chains C and E (**Figure 4C**). Each asymmetric unit contains two epitope footprints, at the i2f and q2f symmetry axes, respectively (**Figure 4C**). At the i2f axis one half of the epitope footprint is located on chain A’ and the other half of on chain A (**Figure 4C**), whereas at the q2f axis, most epitope residues are on chain C, with some on chain E. (**Figure 4C, S4B**). At the i2f axis, one bound Fab will exclude the other i2f site related by two-fold symmetry from being bound. Whereas, at the q2f axis, the position on chain C and E is fully occupied. The result is an occupancy of 1.5 Fab per asymmetric unit.

Cryo-EM density data at the q2f axis identified as the Fab constant and variable domains occupied a single position (**Figure 4A**). However, two distinct positions of Fab constant domain densities could be identified at the i2f axis (**Figure 4A, S4B**). Hence, at the i2f axis the epitope can be approached from two angles but at the q2f it is only approached at one angle. At the i2f axis, only one apparent position for variable domain density was identified. It is most likely an average of two distinct variable domains bound to the virus surface at different constant domain angles (**Figure S4B**).

A top-down view of the virus particle shows that Fab densities at the q2f and i2f axes form a ring around the 5-fold axis of symmetry (i5f) with the constant domains pointing toward the i5f axis (**Figure S4C**). Computational modeling of the DH1017.IgM pentamer (PDB 2CRJ) above the surface of the virion suggests the Fc domain can be centered and positioned above the i5f axis with the Cμ3 ends above the constant domains of the Fabs (**Figure 5A**). The IgM Cμ2 domains are flexible and can bend up to 90 degrees relative to the planar Cμ4-Cμ3-J portion, facilitating a bent, umbrella-like, IgM conformation (Keyt et al., 2020; Murphy et al., 2012; Sharp et al., 2019). The flexibility at Cμ2 domain suggests it is possible the arms of the DH1017.IgM pentamer can bend toward the surface of the virus and each arm can contact the epitopes at q2f and i2f of neighboring asymmetric units (**Figure 5A**). Such arrangement of the epitopes on the virion surface may allow Fab pairs of a pentamer arm to contact epitope pairs on the virus surface either at an i2f and q2f axes between neighboring asymmetric units or at the i2f and q2f axes within the same asymmetric unit simultaneously, allowing DH1017.IgM a decavalent mode of epitope recognition.

**Figure 5:**
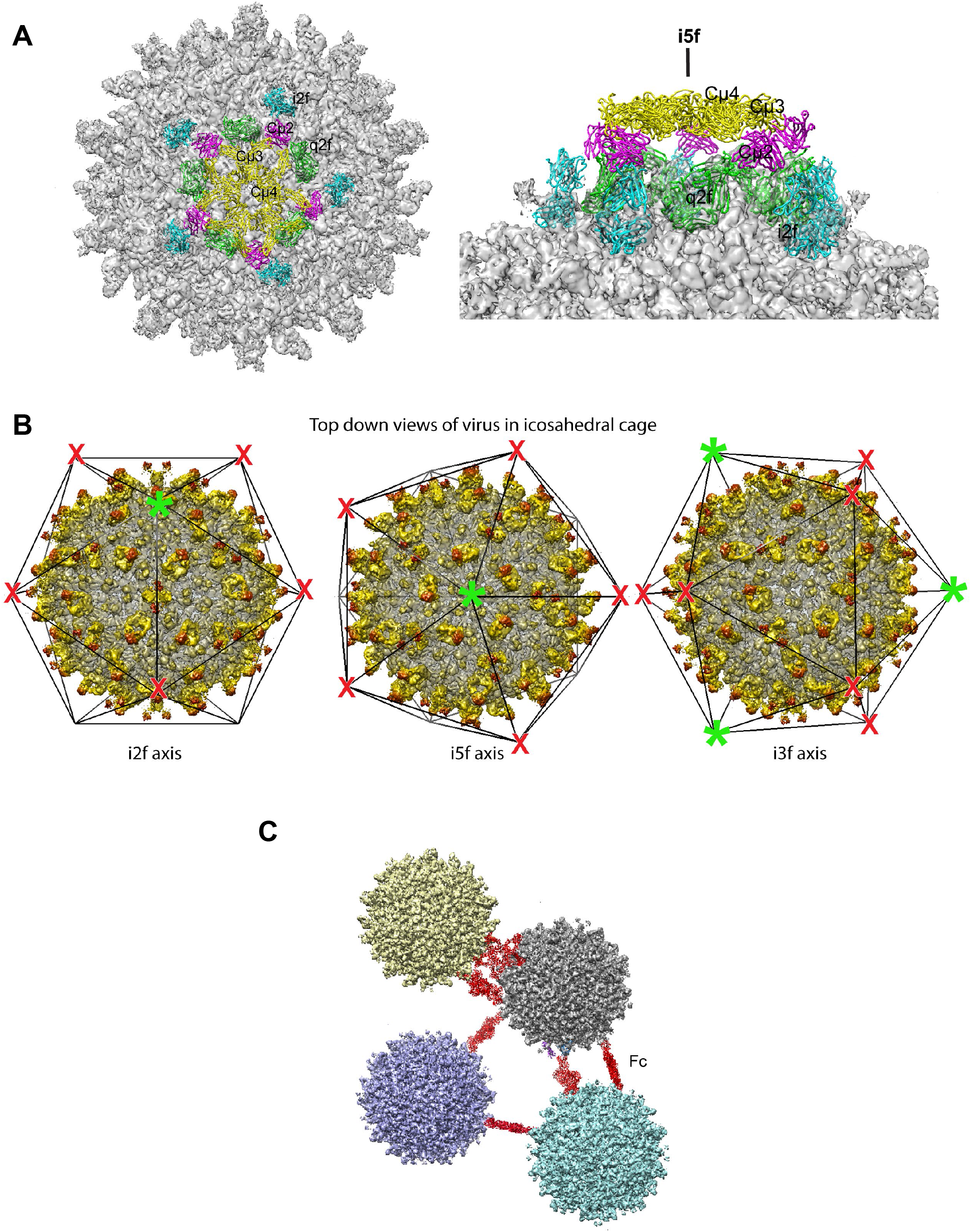
Models of IgM pentamer bound to Zika virus particle. **A.** A top down and side view of a model of the pentamer binding the Zika virus particle in an umbrella like conformation. The density map is shown in gray. The Fc domain is shown in yellow, domains Cμ3 and Cμ4 are labeled. The Cμ2 domain is labeled and colored purple. The DH1017 Fab structure is fit to the density map, at the i2f it is cyan and at the q2f, green. **B.** The density map of the Fab bound virus particle in an icosahedral cage with top-down views at each icosahedral axis, i2f, i5f, and i3f. The green asterisk indicates a position where the pentamer is bound as in A. The red X indicates the position of five-fold axes related to the green asterisk by two-fold symmetry that do not have the i2f position available for Fab binding to form the umbrella like conformation. The i3f view shows positions of complete occupancy for the particle bound as in A. **C.** Schematic representation of virus particles crosslinked by pentamers. The Fc portion of the molecule is shown in red crosslinking virus.

In the umbrella-like conformation, a theoretical maximum of three pentameric IgM can bind concurrently to the same virion, occupying 30 of the 90 epitopes present on the virion surface (**Figure 5B**). In this bent conformation, the IgM pentamer can contact five epitope pairs compared to a single epitope pair for the recombinant DH1017.IgG due to the decavalent versus bivalent mode of antigen recognition. Alternatively, the less bent and planar pentamer conformations may bind one or more pairs of epitopes on one virus particle and bind one or more pairs of epitopes on a second virus particle (**Figure 5C**). This mode of binding effectively cross-links multiple virus particles into aggregates. Simultaneous engagement of five E dimers at the 5-fold axis of viral symmetry demonstrates a novel mechanism of IgM-mediated neutralization of ZIKV that is not available to an IgG molecule and may contribute to the dramatically enhanced IgM potency compared to the IgG.

### DH1017.IgM protects against lethal ZIKV challenge in a mouse model

Finally, we sought to evaluate whether DH1017.IgM could protect against ZIKV infection in mice. We administered 100μg of DH1017.IgM or a non-ZIKV binding human IgM to *Ifnar1*^-/-^ mice 1 day prior and 1 day following infection with 1000 FFU of ZIKV H/PF/2013 and measured viremia and lethality. We found that DH1017.IgM conferred protection from lethal challenge in all mice and controlled viremia to the limit of detection (3.2-3.6 Log_10_ viral copies/mL) in comparison to mice receiving control IgM (6.3-6.9 Log_10_ viral copies/mL). In contrast, all mice in the control IgM group succumbed to infection (**Figure 6A, B**). Human IgM was maintained *in vivo* at detectable levels up to 4 days post challenge in both groups, or 3 days after the last administration (**Figure 6C**). Thus, administration of ultrapotent DH1017.IgM protects against ZIKV disease in mice.

**Figure 6:**
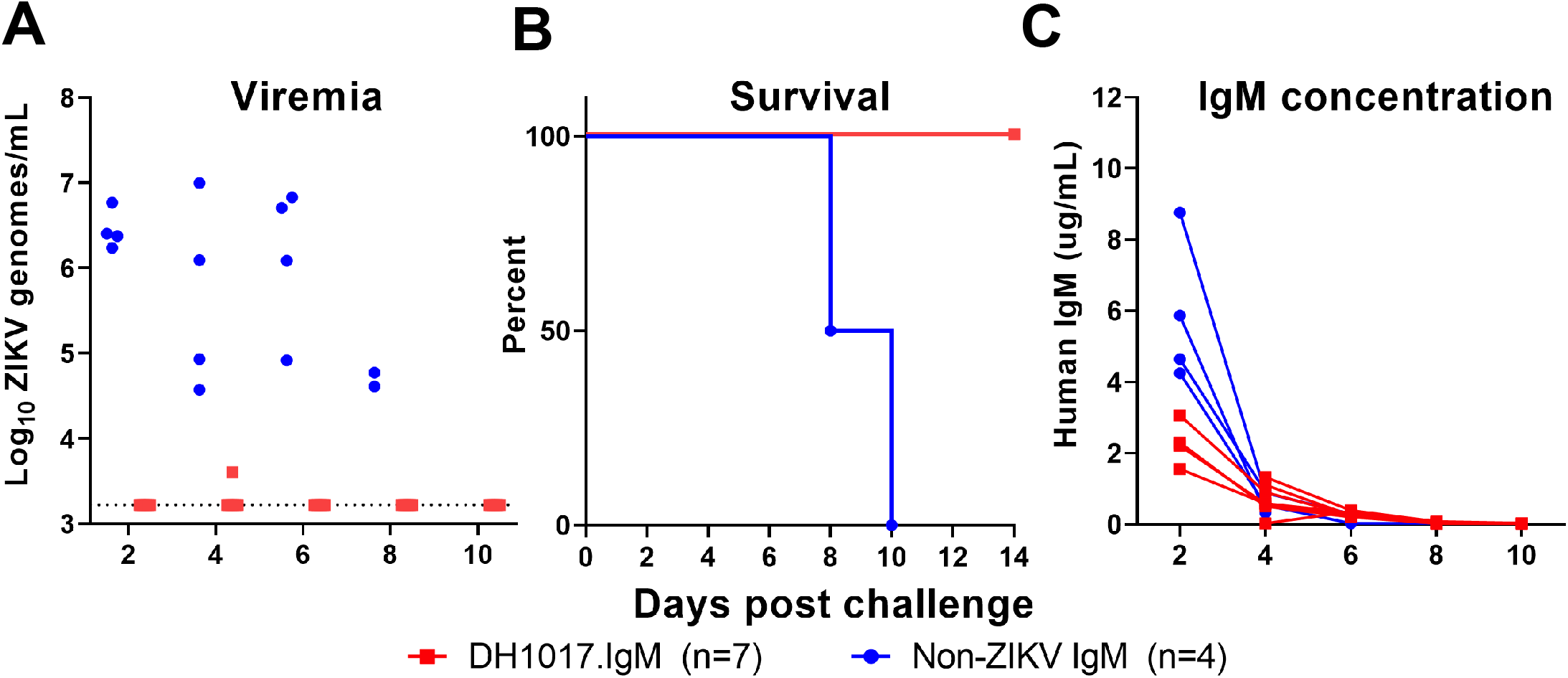
DH1017.IgM protects mice against lethal ZIKV challenge. Five-week-old *Ifnar1*^-/-^ mice were challenged with 1000 focus forming units (FFU) of ZIKV H/PF/2013 in the footpad on day 0. At 1 day prior, and 1 day post challenge, mice were intravenously administered 100μg of human IgM monoclonal antibody, either DH1017.IgM (n=7) or ZIKV non-binding IgM (n=4). Sera was sampled serially up to 14 days post challenge. **A.** Viral load in serum was assessed by qRT-PCR and limit of detection was 1000 copies/mL. **B.** Survival curves for each IgM intervention group. **C.** Total human IgM concentrations were measured in mouse sera by ELISA over days post challenge. Limit of detection for this assay was 0.08-0.03 μg/mL across assays.

## Discussion

In this study, we evaluated the contribution of plasma IgM responses to ZIKV neutralization in pregnancy, defined the frequency of ZIKV-specific B cells isolated from peripheral blood, and isolated a ZIKV-specific IgM monoclonal antibody, DH1017.IgM, in its native multimeric form. Unlike previous plasma-depleted and whole plasma investigation of IgM, the isolation of a human IgM mAb in its native multimeric conformation enabled direct characterization of the impact of the IgM isotype on antibody function against ZIKV. DH1017.IgM did not cross-react with DENV 1-4 and mediated ultrapotent ZIKV neutralization. DH1017.IgM ultrapotency depended on its isotype, as it neutralized >40-fold more potently than when expressed as an IgG. We mapped the footprint of its discontinuous epitope within the asymmetric unit and predicted a mode of antigen recognition compatible with the concurrent engagement of all the ten antigen-binding sites present on the IgM pentamer, a solution not available to smaller IgG molecules with only two antigen-binding sites. These findings identify a unique functional niche of IgM antibodies in protection against ZIKV. Finally, we demonstrated in a murine model that DH1017.IgM protects from ZIKV challenge.

The humoral response to ZIKV infection, like other flaviviruses, is characterized by the development of persistent IgM serum antibodies (Gibney et al., 2012; Griffin et al., 2019; Monath, 1971; Stone et al., 2020). Recent studies have demonstrated that persisting ZIKV-specific serum IgM antibodies predominantly contribute to ZIKV neutralization (Calvert et al., 2021; Malafa et al., 2020). ZIKV infections during pregnancy are of particular concern due to the risk of congenital transmission to the fetus and subsequent burden of disease in children. Pregnancy is characterized by physiological alterations of the immune system but the characteristics of the ZIKV-specific IgM response in pregnant women remained unstudied. In our cohort of pregnant women, circulating IgM antibodies contributed to ZIKV neutralization primarily within the first 3 months of infection and as a minor contributor up to 209 DPS, which aligns with a previous report of ZIKV-specific IgM nAb dynamics outside pregnancy (Calvert et al., 2021). The temporal association (10-14 days post infection) between clearance of ZIKV infection in most individuals (Lessler et al., 2016; Paz-Bailey et al., 2018) and the early peak of serum IgM which precedes peak IgG responses (Ravichandran et al., 2019; Tonnerre et al., 2020) suggests that early IgM neutralizing responses may play an important role in early control of infection. Our data extend such temporal association in the setting of pregnancy.

We note that subject P73, from whom we isolated DH1017.IgM at day 71 DPS, developed a prolonged viremia lasting 42 days despite an early robust peak in plasma IgM neutralization. However, while viremia outlasted the first plasma IgM neutralization peak in this subject, high levels of IgM ZIKV neutralization persisted up to 71 DPS, 30 days after viremia control. The relationship between prolonged viremia and robustness of plasma IgM should be further investigated.

While the role of serum IgM in flavivirus infection is actively being studied, ZIKV-specific IgM monoclonal antibodies have not been characterized, hindering our understanding of the mechanisms of action of this isotype against ZIKV. From subject P73, we derived a fully human IgM-expressing B-LCL secreting the ZIKV-specific ultrapotent DH1017.IgM nAb in its native multimeric conformation. While subject P73 had a prior exposure to DENV, DH1017.IgM did not bind to DENV and was therefore not part of a recall response from pre-existing DENV immunity. DH1017.IgM was nonetheless somatically mutated. The original B cell was sorted from unfractionated B cells including naïve, class-switched, and non-class-switched memory B cells.

Naïve B cell can undergo SHM *in vitro* upon EBV transformation after prolonged culturing (≥28 days), often with evidence of clonal diversification, and can class-switch when stimulated with CD40L and IL-21 (Heath et al., 2012), which were included in our stimulation regimen. We did not identify additional nucleotide mutations or class-switched DH1017.IgG mAb from the DH1017.IgM LCL throughout multiple rounds of prolonged cell expansions. Hence, while we cannot exclude an origin from a naïve B cell, these observations suggest that DH1017.IgM LCL was derived from an IgM+ memory B cell.

Structural analysis of DH1017.Fab in complex with whole ZIKV virion identified a quaternary epitope footprint encompassing primarily DII that differs from that of known potently ZIKV-neutralizing mAbs. Antibody responses targeting quaternary epitopes on multimeric E protein epitopes or mature virions are associated with protection from ZIKV infection in mice (Maciejewski et al., 2020; Metz et al., 2019). Recognition of a quaternary epitope on ZIKV is the defining feature of a class of potently neutralizing mAbs (Collins et al., 2019; Hasan et al., 2017b; Long et al., 2019; Rogers et al., 2017; Sapparapu et al., 2016; Stettler et al., 2016).

Upon viral entry into cellular endosomes, E dimers rearrange into fusogenic trimers to mediate membrane fusion (Smit et al., 2011) and DH1017.IgM binding across Domain II of the E dimer may cross-link ZIKV E monomers within a single dimer and restrict conformational changes to prevent infection (Zhang et al., 2015). Such mechanism has also been proposed for ZIKV-117 and ZIKV-195 IgG mAbs (Hasan et al., 2017b; Long et al., 2019). However, our data demonstrate that IgM ultrapotent neutralization is largely depended on its isotype, implying additional non-mutually exclusive modes of isotype-dependent neutralization that are not available to IgG.

First, an IgM may efficiently cross-link multiple virions into aggregates that would not efficiently internalize into cells via endosomes. Second, the large molecular size of an IgM may interfere more efficiently than an IgG with virion attachment to the cell membrane and prevent cell entry. Third, IgM may bind with high orders of valency to a virion by virtue of its multimeric structure, thus cross-linking multiple E dimers. This could make the particle more rigid and sterically hinder interactions with the cell surface to prevent either binding or fusion with the host cell. Based on the position of the DH1017.IgM epitope on the i2f and q2f axes of symmetry within each asymmetric unit, computational modeling of the DH1017.IgM pentamer predicts cross-linking of five E dimers at the five-fold axis of symmetry resulting in a decavalent mode of antigen recognition. However, concurrent cross-linking of multiple virions and competition with other ZIKV-binding antibodies *in vivo* may interfere with a decavalent mode of antigen recognition on a single virion. Conversely, the pentameric IgM may out compete other less avid IgG mAbs.

Functional results together with the potential for a multivalent mode of antigen recognition suggests a functional niche that is exclusive to IgM in the context of pathogens with repetitive proximal structures. The arrangement of the cognate epitope relative to the five-fold axis of symmetry on the virion surface is essential for multivalent antibody interactions. Whether neutralizing epitopes other than the DH1017 epitope are permissive to multivalent interactions needs to be determined.

Intriguingly, class-switching to IgG of the DH1017.IgM clone would result in reduced function. Thus, we speculate that the non-class switched IgM+ memory B cell and IgM+ plasma cell pools could provide a source of neutralizing antibodies that interact with virions in an isotype-exclusive manner. Future studies should address if the IgM+ memory B cell compartment is a preferential source of ZIKV IgM neutralizing antibodies.

Neutralizing antibodies mediate protection against ZIKV as demonstrated in transfer of passive immunity via immune plasma in the non-human primate model (Larocca et al., 2016; Richner et al., 2017). However, IgG neutralizing antibodies at sub-neutralizing concentrations can enhance flavivirus replication via ADE through FcγR mediated endocytosis of the immune complex. *In vitro*, DH1017.IgG mediated ADE, whereas DH1017.IgM did not. We note that these experimental results may be influenced by differences in the relative expression of FcγR and FcμR/Fcα/μR on cell surface. In addition, DH1017.IgM *in vitro* neutralization potency increased in presence of complement in a dose-dependent manner. Since this complement-mediated effect lowers the stoichiometry requirements for neutralization, as previously reported for West Nile virus (Mehlhop et al., 2009), and considering the high potency of DH1017.IgM, the risk of DH1017.IgM-mediated ADE *in vivo* may be further reduced. Importantly, DH1017.IgM mediated protection against subcutaneously administered lethal ZIKV challenge in mice, recapitulating protection and viral control conferred by potent IgG neutralizing antibodies previously tested (Collins et al., 2019; Long et al., 2019; Robbiani et al., 2017; Sapparapu et al., 2016; Swanstrom et al., 2016).

Since efficacy trials for ZIKV vaccines are limited by currently low levels of endemic circulation, the development of prophylactic interventions that can be safely deployed to pregnant women during the next ZIKV outbreak are urgently needed to mitigate the risks of congenital ZIKV transmission. We propose that DH1017.IgM may be suitable as a passive intervention for protection against ZIKV infection. In particular, since IgM isotype antibodies are not transferred across the placenta, the potential risks of fetal toxicity and ADE in infancy after transplacental IgG transfer are expected to be mitigated. Thus far, 20 IgM mAb interventions have been tested in clinical trials and demonstrate that infused IgM are safe and well tolerated (Keyt et al., 2020).

Since the IgM half-life is considerably shorter than IgG, antibody engineering to prolong the half-life would be likely required for effective prophylactic countermeasures. However, the potential issue of short half-life would be less relevant for a therapeutic intervention administered at the time of diagnosis, which will be aimed at rapid clearance of viremia and reduction in the time of fetal exposure to circulating virus. Further, our findings support the development and investigation of engineered multimeric antibody formulations as a prophylactic and therapeutic strategy.

In summary, we demonstrated a large contribution of plasma IgM in ZIKV neutralization in both primary and DENV pre-exposed ZIKV infections in pregnancy. We isolated a novel ultrapotent ZIKV-neutralizing IgM mAb that protected mice from lethal challenge and demonstrated the impact of isotype on its antiviral function. We defined a conceptual framework by which the spatial arrangement of quaternary epitopes on the virion can modulate neutralization in an isotype-specific manner. As congenital transmission of ZIKV in pregnancy is the source of the greatest burden of ZIKV disease, further studies are warranted to assess whether DH1017.IgM can protect against fetal infection in pregnant animal models and mitigate fetal damage (Van Rompay et al., 2020) by reducing or preventing maternal viremia. With the experience of delayed roll out of vaccines to pregnant women for emerging outbreaks such as Ebola and more recently the SARS-CoV-2 pandemic (Craig et al., 2021; Gomes et al., 2017), a prophylactic intervention that is rapid, protective, and safe for use pregnancy will be particularly needed when ZIKV re-emerges.

## Acknowledgments

This work was supported by a Duke Global Health Institute Award to S.R.P. and a Duke Incubation Fund Award to M.B.; by NIH/NIAID grants R21-AI132677 (S.R.P.), R21-AI144631 (H.M.L.), F31-AI143237 (C.A.L.) and R01-AI073755 (R.J.K., PI: M.S. Diamond), by NIH/NIAID contract HHSN272201700060C (R.J.K; PI: K. Satchell) and in part by the Division of Intramural Research, NIAID, NIH (M.B.). M.B. was employed by the Duke Human Vaccine Institute (DHVI) at the time of this work and is currently employed by the Division of Intramural Research, NIAID, NIH. C.G. was supported by the Coordenação de Aperfeiçoamento de Pessoal de Nível Superior - Brasil (Finance Code 001). The pregnancy cohort in Brazil was funded by the Fundação de Amparo à Pesquisa do Espírito Santo (306/2016 to RD; Protocol Financial Support Number: 74910132/16). We thank Drs. Aravinda de Silva (UNC Chapel Hill) and James Crowe (Vanderbilt University) for sharing reagents; Madison Berry, Drs. M. Anthony Moody and Kevin Wiehe (DHVI), Brian Watts of the DHVI BIA Core Facility and the DHVI Research Flow Cytometry Facility for helpful scientific discussions and/or logistical and technical support. Also, we thank Juliana Carnielli, Solange Alves Vinhas and Keyla Fonseca for establishing cohort enrollment and sample collections at Federal University of Espírito Santo. We are grateful to the maternity ward of Cassiano Antonio de Moraes Hospital staff for supporting the participants’ deliveries and birth sample collections. Importantly, we thank the mothers who participated in this study and made efforts to provide samples for research during pregnancy.

## Contributions

Conceptualization, T.S., R.D., S.R.P. and M.B.; Methodology, T.S., A.S.M., C.A.L., L.P., R.D., H.M.L., R.J.K., S.R.P. and M.B.; Investigation, T.S., K.K.H., A.S.M., R.L.J., C.A.L., M.A.G., I.M., H.S.W., J.A.E., K.L., T.V.H., R.J.E., S.V., K.E.B., S.Z., J.F.M., J.J.T., M.D., L.P. and M.B.; Resources, C.G., L.P., R.D., T.C.P., E.E.O., H.M.L., S.R.P. and M.B.; Writing – Original Draft, T.S., A.S.M., S.R.P. and M.B.; Writing – Review & Editing, all authors; Visualization, T.S., A.S.M., C.A.L. and M.B.; Supervision, T.C.P., E.E.O., H.M.L., R.J.K., S.R.P. and M.B.; Funding Acquisition, C.A.L., C.G., R.D., H.A.L., R.J.K., S.R.P and M.B.

## Methods

### Donors and Sample Information

Participants in this study were enrolled from July 2016 to October 2017 in the city of Vitória, State of Espírito Santo, in Brazil. In this prospective cohort, we enrolled pregnant women with Zika-like symptoms of rash and/or fever and collected blood samples through pregnancy, delivery, and postpartum. Cohort design, recruitment, enrollment and sample collection are detailed in our prior study (Singh et al., 2019). Plasma and peripheral blood mononuclear cells (PBMCs) from 11 mothers who were serologically confirmed for ZIKV infection previously (Singh et al., 2019) were included in this study. The identifiers of the subjects included in this study are as follows (alternative identifiers in parenthesis): P14 (B1_0001), P15 (B1_0002), P17 (B1_0004), P19 (B1_0005), P23 (B1_0007), P24 (B1_0008), P34 (B1_0014), P50 (B1_0027), P54 (B1_0027), P56 (B1_0031) and P73 (B1_0037). The infant born to subject P14 demonstrated microcephaly at birth, and the rest had no known signs of congenital Zika syndrome. DH1017.IgM was isolated from the PBMCs of P73, who presented with a symptomatic ZIKV infection at 22 weeks of gestation and had prolonged viral replication with vRNA detected in serum/urine up to 42 DPS. She delivered an apparently healthy baby at 38.5 weeks of gestation with normal head circumference. All other mothers had detectable ZIKV vRNA in plasma at the first visit only.

### Ethics Statement

The Institutional Review Boards of Hospital Cassiano Antonio Moraes (Brazilian National Research Ethics Committee (CEP/CONEP) Registration number: 52841716.0.0000.5071) and the Duke University Medical Center (Pro00100218) each approved this prospective cohort study and sampling. Pregnant women with rash or fever, who were a minimum of 18 years of age, and provided a willingness to participate through a written informed consent were approved for inclusion into this study. To protect the privacy of all subjects, publicly available identifiers are twice removed from the participant. All practices conducted as part of this study are compliant with ethical principles for medical research involving human subjects as outlined in the Declaration of Helsinki.

Animal work was approved by the University of North Carolina at Chapel Hill IACUC (protocol no. 20-132).

### Cell culture and virus stocks

Vero-81 cells were grown in Dulbecco’s Modified Eagle Media (Gibco 11965092) supplemented with 10% heat-inactivated fetal bovine serum (Sigma, F4135-500mL), 1x penicillin-streptomycin (Gibco 15140-122), and 1x MEM non-essential amino acids solution (Gibco 11140-050). Viruses used for the focus reduction neutralization test were DENV1 (WestPac74), DENV2 (S-16803), DENV3 (CH54389), DENV4 (TVP-360), provided by Dr. Aravinda de Silva (University of North Carolina at Chapel Hill); ZIKV H/PF/2013 and PRVABC59 strains were obtained from the Biodefense and Emerging Infections Research Resources Repository (BEI). Virion binding antibodies were detected using the following viruses from BEI: ZIKV (PRVABC59), DENV1 (Hawaii NR-82), DENV2 (New Guinea C), DENV3 (Philippines), and DENV4 (H241). ZIKV was grown in Vero-81 cells supplemented with 10% heat-inactivated fetal bovine serum (FBS) and 10mM HEPES (Sigma H0887-100ML). Dengue viruses were grown in C6/36 cells cultured in RPMI 1640 (Gibco 11875-093), L-Glutamine (Gibco 25030-081), 25 mM HEPES (Gibco, 22400-089), 1x penicillin-streptomycin (Gibco, 15140-122), and 20% FBS.

During DENV infection, RPMI was supplemented with 25mM HEPES and 2% FBS. 0.5 mL of viruses were added to 80-90% confluent cells. Cells were infected with DENV2, DENV3, or DENV4 for 7-9 days and DENV1 for 11 days. ZIKV stock infections were stopped when cytopathic effect was observed (∼3-6 day infection). Cell supernatant containing virus was then harvested, centrifuged brought to a final concentration of 20% FBS, and filtered through 0.22µm filter prior to storage at −80°C for use. FcγR-expressing K562 cells and THP-1 cells were maintained in RPMI 1640 medium containing Glutamax (Invitrogen) with 7% heat-inactivated FBS (Invitrogen) and 1X penicillin-streptomycin (Invitrogen).

### Fluorescent labelling of ZIKV for sorting B cells

A previously developed approach to label DENV with Alexa Fluor 488 (AF488) dye was adapted to label ZIKV (Zhang et al., 2010). ZIKV (Strain: PF13/251013-18) was propagated in Vero cells and purified through 30% sucrose. Virus titer was determined using the BHK-21 cell plaque forming assay as previously described (Chan et al., 2011). Briefly, AF488 succinimidyl ester was reconstituted in 0.2M sodium bicarbonate buffer (pH 8.3) and added to 3 x 10^8^ PFU of ZIKV at a final dye concentration of 100 µM. The mixture was incubated at room temperature for 1 hour with gentle inversions every 15 minutes. Labelled ZIKV was purified by size exclusion chromatography using Sephadex G-25 columns (Amersham, GE Healthcare) to remove the excess dye and titrated again on BHK-21 cells. Thereafter, ZIKV was UV inactivated (254nm) for 1 minute on ice. Inactivation of virus was verified by serial passaging of virus on C6/36 mosquito cell line (ATCC), without detection of infection on this susceptible cell line.

### Staining and sorting ZIKV-reactive B cells

Thawed PBMCs were stained with Aqua Vital Dye (Invitrogen), IgD-PE (clone IA6-2; BD Biosciences), CD10 PE-CF594 (clone HI10a; BD Biosciences; CD3 PE-Cy5 (clone HIT3a; BD Biosciences), CD14 BV605 (clone M5E2; BD Biolegend), CD16 BV570 (clone 3G8; Biolegend), CD27 PE-Cy7 (clone O323; Thermo Fisher Scientific), CD38 APC-AF700 (clone LS198-4-3; Beckman Coulter), CD19 APC-Cy7 (clone SJ25C1; BD Biosciences) and 1 x 10^6^ PFU of freshly thawed UV-inactivated ZIKV labelled with AF488. Additionally, 5µM of Chk2 kinase inhibitor II Inhibitor (Calbiochem/EMD Chemicals) was added to prevent cell death. Unfractionated B cells were defined as CD14^-^/CD16^-^/CD3^-^/CD19^+^. For memory B cells an additional IgD^-^/CD27^all^ gate was applied. The ZIKV-reactive AF488 gate was set using a fluorescence minus one condition. Cells were sorted on a Beckton Dickinson FACS Aria II and analysis was performed in FlowJo.

### B cell cultures

ZIKV-reactive B cells were cultured as previously described (Bonsignori et al., 2011, 2016). Briefly, cells were sorted in wells pre-seeded with human CD40 ligand-expressing MS40L cells (5,000 cells/well) and containing 2.5µg/ml CpG ODN2006 (tlrl-2006; InvivoGen), 5µM CHK2 kinase inhibitor II) and 50 ng/ml of recombinant human (rHu) IL-21 (200-21; Peprotech) in RPMI/15% fetal calf serum. EBV was added for the overnight incubation (200µl supernatant of B95-8 cells per 10^4^ B cells). After overnight incubation in bulk at 37°C in 5% CO_2_, B cells were distributed by limiting dilution at a calculated concentration of 1 cell/well into 96-well round-bottom tissue culture plates in the presence of MS40L cells and complete medium described above. Medium was refreshed after 7 days. Supernatants were collected after 14 days. Cell culture supernatants were assessed for IgG, IgM, and IgA levels in ELISA as previously described (Bonsignori et al., 2016) and Ig+ wells were further evaluated for ZIKV-reactivity with a virion binding ELISA. From each culture well, half of the cells were harvested and preserved in RNAlater (Qiagen) at day 14 and half were maintained in culture. Cell clones that immortalized were expanded and cloned by using a standard limiting dilution method. Reactivity and IgV(D)J sequences were checked periodically. We derived three IgG+ B-LCLs from P34 (termed 119-4-D6.IgG, 119-1-D7.IgG, and 119-5-C5.IgG), three IgG+ B-LCL from P56 (124-4-C8.IgG, 124-1-C2.IgG and 124-2-G3.IgG), and one IgG+ B-LCL from P54 (126-1-D11.IgG), From subject P73 we derived two B-LCLs: one expressing IgG from memory B cells collected 28 DPS (123-3-G2.IgG), and one IgM+ B-LCL (termed DH1017) from unfractionated B cells collected at 71 DPS.

### Isolation of V(D)J immunoglobulin regions

RNA from positive cultures was extracted by using standard procedures (RNeasy minikit; QIAGEN), and the genes encoding Ig V_H_DJ_H_ and V_L_J_L_ rearrangements were amplified by RT and nested PCR without cloning by use of a previously reported method (Liao et al., 2009). Ig V(D)J sequences (Genewiz) were assembled using a customized bioinformatic tool developed by Kevin Wiehe (DHVI) based on the Cloanalyst software package (http://www.bu.edu/computationalimmunology/research/software/).

### Monoclonal antibody production

B-LCLs were expanded in CELLine bioreactor flasks (Wheaton) following the manufacturer’s recommendations. Monoclonal antibodies were purified using protein A resin columns for IgG and CaptureSelect beads (Thermo Scientific) for DH1017.IgM following manufacturer’s recommendations. DH1017.IgG and DH1017.Fab heavy and light chain plasmids (GenScript) were co-transfected at an equal ratio in suspension Expi 293i cells (Invitrogen) using ExpiFectamine 293 transfection reagents (Life Technologies) according to the manufacturer’s protocols. Transfected cells were gently shaken overnight for 16-18 hours and incubated at 37°C with 8% CO_2_. We then added the enhancer provided in the kit and incubated at 37°C with 8% CO_2_ for 4-6 days. Supernatant containing antibody was harvested and filtered, and co-incubated with a Protein A affinity resin (Thermo Fisher Scientific) for IgG antibody or LambdaFabSelect Agarose Beads (GE Healthcare Life Sciences) for Fab at 4°C on a rotating shaker overnight. The bead and supernatant mixture was then loaded onto a column for purification. Following a Tris/NaCl Buffer (pH 7.0) wash, mAb was eluted from beads using Trizma HCl (pH 8.0; VWR) and concentrated in the Vivaspin Turbo 15 Concentrator (Thermo Fisher Scientific) with a pH neutralizing buffer exchange using Citrate Buffer (pH 6.0). Purified antibody concentration was determined by Nanodrop and product was evaluated by reducing and nonreducing SDS-polyacrylamide gel electrophoresis and Coomassie Blue staining for appropriate size.

### SDS PAGE and Coomassie

DH1017.IgM was run under non-reducing conditions using a NuPAGE 3-8% Tris-Acetate Gel (Invitrogen) with 1x Tris-Glycine Native Running Buffer (Novex) at 130V for 2.5 hours. DH1017.IgM (5µg/lane) was prepared with Native Tris-Glycine Sample Buffer (Novex) and NativeMark Protein Standard (Invitrogen) was used as the ladder. Gel was subsequently stained with Coomassie and imaged using ChemiDoc (Bio-Rad).

### Negative stain electron microscopy

An aliquot of DH1017.IgM was equilibrated to room temperature, then diluted to 20-40 µg/ml with buffer containing 10 mM HEPES, pH 7.4, 150 mM NaCl, and 0.02% ruthenium red. A 5 µl drop of diluted sample was applied to a glow-discharged carbon-coated grid for 8-10 seconds, blotted, then rinsed with two drops of buffer containing 1 mM HEPES, pH 7.4 and 7.5 mM NaCl, and finally stained with one drop of 2% uranyl formate for 60 s, then blotted and air dried. Images were obtained with a Philips EM420 electron microscope at 120 kV, 82,000x magnification, and captured with a 2k x 2k CCD camera at 4.02 Å/pixel. The RELION program (Scheres, 2016) was used for particle picking and 2D class averaging.

### ZIKV and DENV virion capture ELISA

The virion capture ELISA methods were previously described (Singh et al., 2019). Briefly, high-binding 96-well plates (Greiner) were coated with 40 ng/well of 4G2 antibody (clone D1-4G2-4-15) in 0.1 M carbonate buffer, pH 9.6 overnight at 4°C. Plates were blocked in Tris-buffered saline containing 0.05% Tween-20 and 5% normal goat serum for 1 hour at 37°C, followed by an incubation with either ZIKV (PRVABC59), DENV1 (Hawaii), DENV2 (New Guinea C), DENV3 (Philippines), and DENV4 (H241) from BEI for 1 hour at 37°C. ZIKV and DENV2 were diluted 1:5, DENV1 and DENV3 diluted 1:3, and DENV4 diluted 1:7. 50 µL/well samples were added and incubated for 1 hour at 37°C. Culture supernatants were screened undiluted whereas plasma and purified mAb binding was measured in duplicate. Eight-point serial dilutions for plasma started at 1:25 with 3-, or 5-fold dilutions and mAbs started at 100 ug/mL 5-fold serial dilutions. Horseradish peroxidase (HRP)-conjugated goat anti-human IgG antibody (Jackson ImmunoResearch Laboratories, 109-035-008), HRP-conjugated goat anti-human IgM antibody (Jackson ImmunoResearch Laboratories, 109-035-129), or HRP-conjugated goat anti-human IgA antibody (Jackson ImmunoResearch Laboratories, 109-035-011) were all used at a 1:5,000 dilution followed by the addition of SureBlue Reserve TMB substrate (KPL, Gaithersburg, MD). Reactions were stopped by Stop Solution (KPL) after five minutes and optical density (OD) was detected at 450 nm (Perkin Elmer, Victor). For IgG and IgM, an isotype matched known ZIKV-specific commercial mAb was used as a positive control (IgG: C10, Absolute Antibody, Cat #: AB00677-10.0; IgM: Anti NS1 IgM, myBiosource, Cat#: MBS6120634). Plasma from ZIKV-infected individuals was used as positive control for IgA culture supernatants. The negative control condition was sample diluent alone. Respiratory syncytial virus specific IgG Palivizumab was used as a negative control for testing ZIKV-binding of purified mAbs, and diluent served as negative control for IgM purified mAb assays. For purified mAbs that were serially diluted, the magnitude of virion binding was evaluated as an ED_50_, which was calculated with a sigmoidal dose-response (variable slope) curve in Prism 7 (GraphPad) using a least squares fit. The ED_50_ value for serially diluted mAbs was considered valid if the OD_450_ at 100ug/mL was 5-fold higher than the no sample condition. For plasma, the ED_50_ value was considered valid if the OD_450_ at 1:25 dilution 2 standard deviations above the mean OD_450_ for DENV 1-4, and 3 standard deviations above the mean OD_450_ observed for 11 plasma samples from healthy U.S. subjects (2SD OD_450_ cut-offs: DENV-1 = 0.406, DENV-2 = 0.648, DENV3 = 0.906, and DENV-4 = 0.885; 3SD OD_450_ cut-off: ZIKV = 0.596). For culture supernatants, the cut-off for positivity was the highest of mean plate blanks OD_450_ + 2 standard deviations and OD_450_ not less than 0.4.

### Recombinant E protein ELISA

DH1017 mAbs was immobilized to a high-binding 96-well plate (Greiner) at 0.5ug/mL in 1x TBS, then blocked with 3% milk in TBS tween, each for 1 hour at 37°C. Serial dilutions of his-tagged recombinant E protein dimer (Premkumar et al., 2017) starting at 20ug/mL with 2-fold dilutions to 12-spots were added to the plate for an hour at 37°C. This was followed by 0.5ug/mL anti-his HRP (Sigma), and binding was detected after incubation with substrate at an absorbance of 450nm. EC_50_ was obtained by a sigmoidal dose-response (variable slope) curve in Prism 7 (GraphPad) using least squares fit. The negative control was no antigen and positive control were previously reported mAbs, including 1M7, ZV-2, ZV-64 and ZKA190. 1M7 mAb (Smith et al., 2013) was produced from a hybridoma. ZKA190 was generated recombinantly using the sequence for PDB entry 5Y0A (Wang et al., 2017). The mouse mAbs ZV-2 and ZV-64 (Zhao et al., 2016) were produced from hybridomas at the UNC protein expression core facility. ZIKA-752, ZIKA-893, and rZIKV-195 were produced and provided by James Crowe.

### Focus reduction neutralization test

We used previously described methods for FRNT_50_ in a 96-well plate (Singh et al., 2019). Briefly, serial dilutions of heat-inactivated plasma or purified mAbs were added to 30-60 focus forming units of either DENV serotypes 1-4 or ZIKV (H/PF/2013). Plasma was used at a starting dilution of 1:25 with subsequent 5-fold or 7-fold dilutions. MAbs were tested at 5ug/mL or 10ug/mL with 5-fold dilution series. However, DH1017.Fab, was tested at 1mg/mL with a 5-fold dilution series. Negative control was media alone, and positive controls were known ZIKV-neutralizing mAbs and plasma from ZIKV-infected subjects. Virus and plasma or mAb samples were co-incubated for 1 hour at 37°C, then transferred to a 96-well plate (Greiner Bio One) with confluent Vero-81 cells and incubated for 1 hour at 37°C. Plates were overlayed with 1% methylcellulose and incubated at 37°C for 40-42 hours (ZIKV and DENV4), 51-53 hours (DENV1), or 48 hours (DENV2 and DENV3). Cells were fixed with 2% paraformaldehyde for 30 minutes and stained with 0.5 μg/mL of 4G2 or E60 mouse monoclonal antibody. Foci were detected with an anti-mouse IgG conjugated to horseradish peroxidase at a 1:5000 dilution (Sigma), followed by True Blue substrate (KPL). Foci were counted using the CTL ImmunoSpot plate reader (Cellular Technology Limited). FRNT_50_ values were calculated with the sigmoidal dose-response (variable slope) curve in Prism 8.3.0 (GraphPad), constraining values between 0 and 100% relative infection. Percent relative infection curves were considered to pass quality control criteria for FRNT_50_ determination if R^2^> 0.65, absolute value of hill slope >0.5, and curve crossed 50% relative infection within the range of the plasma dilutions in the assay. Samples were repeated up to 3 times to quantify a valid FRNT_50_ in accordance with the quality control criteria.

### Reporter Virus Particle (RVP) production

ZIKV RVPs incorporating the structural proteins (Capsid [C], premembrane [prM], and E) of ZIKV strain H/PF/2013 were produced by genetic complementation of a green fluorescent protein (GFP)-expressing WNV lineage II sub-genomic replicon with the virus structural gene plasmids as previously described (Dowd et al., 2016). Briefly, the WNV replicon and C-prM-E plasmids were co-transfected into HEK-293T cells using Lipofectamine 3000^TM^ transfection reagent (Invitrogen). Transfected cells were incubated at 30°C and RVP-containing supernatants were harvested 3-6 days post-transfection and pooled. RVP-containing supernatants were passed through a 0.2 μm filter (Millipore) and stored at −80°C until use.

### RVP-based Antibody-Dependent Enhancement (ADE) assay

RVP-based ADE assays were performed by incubating GFP-encoding RVPs with serial dilutions of mAbs for 1 h at 37°C. FcγR+ K562 or THP-1 cells were infected with RVP immune complexes and incubated at 37°C for 36-48 h. Cells were fixed with paraformaldehyde, and GFP expression was detected by flow cytometry. Antibody enhanced infection was scored as detectable if the number of GFP positive cells was ≥3-fold above background (defined as the average percent GFP positive cells in the absence of antibody).

### Plaque assay-based ADE assay

DH1017.IgM and DH1017.IgG were serially diluted 2-fold over 6.25 – 0.003 µg/mL and 10uL of diluted mAb were co-incubated in duplicate with 2 x 10^5^ PFU of ZIKV (Strain: PD13/251013-18) in a round bottom 96 well-plate for 1 hour at 37°C to form immune complexes. Thereafter, 2 x 10^4^ cells of THP1.2S or primary monocytes were added to the ZIKV and mAb immune complexes for a 72-hour multiday infection at 37°C. THP1.2S cells are a subclone of the THP1 monocyte cell line (ATCC) that are shown to be more susceptible to flavivirus infection (Chan et al., 2014). Collection and processing of primary monocytes has been previously described and was conducted approval of the NUS Institutional Review Board under reference code B-15-227 (Chan et al., 2019). After the multiday infection on monocytes, supernatant was collected, and infectious virus was tittered via plaque forming assays on BHK-1 cells (ATCC) in 24 well plates in quadruplicate, as previously described (Chan et al., 2011). For the positive control we used humanized IgG1 4G2 mAb, and for the negative control we used 3H5 (DENV-specific mAb) as well as the virus only condition, without any mAb present.

### Antibody-dependent complement antiviral activity

Focus reduction neutralization test (FRNT) in the presence of increasing concentrations [volume/volume] of complement from normal human serum (Sigma), including 0, 5, 10, 15, 20 and 25% final concentration were tested. Percent relative infection was calculated as a ratio of the foci in the plasma, ZIKV (PRVABC59), and complement condition, to the foci in the virus and complement condition, multiplied by 100. This approach allows for determination of the antibody-dependent complement activity and adjusts for antibody-independent complement activity. Samples were run in triplicate and experiment was independently repeated for each concentration of complement. Positive control was DH1017.IgM and DH1017.IgG run in the absence of complement, and negative control was the virus and complement only conditions. The rest of the analysis and approach are identical to the FRNT approach.

### Determining the contribution of plasma IgM to ZIKV neutralization

Each original plasma sample was split into 2 aliquots, with one portion depleted of IgM isotype antibodies and another portion mock depleted. First, each sample was heat inactivated for 30 minutes at 56°C, diluted 1:1 with sterile PBS, and centrifuged at 10,000 x G for 10 minutes to remove debris. Depletion beads were packed into sterile 0.5mL centrifugal filter devices with a 0.22µm pore PVDF membrane (Millipore) and equilibrated with 3x sterile PBS washes (pH 7.2). 200mg of POROS^TM^ CaptureSelect^TM^ IgM affinity beads (ThermoFisher Scientific) were used for IgM depletion, and 66mg of corresponding beads of the same size (200-400 mesh) and material (polystyrene divinylbenzene 1% cross linked beads; Alfa Aesar) were used for mock depletion. Samples were co-incubated with beads for 10 minutes at room temperature with gentle inversions, and then the depleted fraction was centrifuged at 10,000 x *g* for 10 minutes.

Depletion of IgM was confirmed by total IgM ELISA, and non-specific losses to ZIKV-binding IgG were quantified through virion binding ELISA; both assays are described in other sections of this Method supplement. Limit of detection (LOD) of 0.12 μg/mL was based on the linear range of a sigmoidal standard curve of human IgM (Jackson ImmunoResearch Laboratories). Magnitude of ZIKV-binding IgG was assessed by virion-binding ELISA and neutralization potency was assessed using the Focus Reduction Neutralization Test. Due to slight differences in ZIKV-binding IgG across IgM and mock depleted fractions, each fraction was adjusted to the magnitude of ZIKV-binding IgG in the same sample such that differences in neutralization activity could be attributed to differences in IgM isotype antibodies. Thus, the percent neutralization attributable to IgM that is reported in this study was calculated as follows:

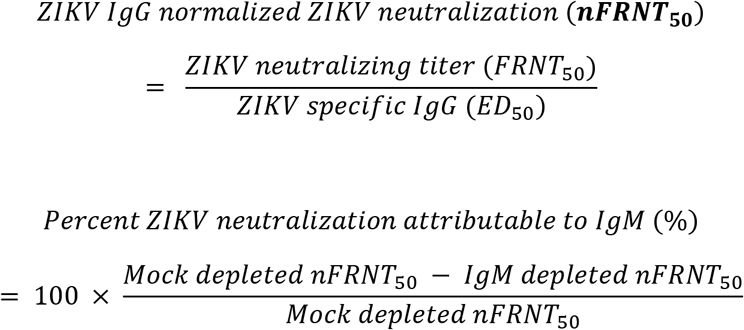

### Detection of antibody isotype from sera

Human IgM antibodies were detected in sera from passively transferred mice using high-binding 384-well plates (VWR) coated with 0.5 µg/well of goat anti-human Ig polyvalent antibody (ThermoFisher, #H17000) in 0.1 M carbonate buffer, pH 9.6 overnight at 4°C. Plates were blocked the next day (2 hours at RT) with 10X PBS containing 4% whey, 15% normal goat serum, and 0.5% tween. Mouse serum samples were diluted 1:30, serially diluted three-fold, then added to the plate (10µL/well), and incubated for 1 hour at 37°C. Horseradish peroxidase (HRP)-conjugated goat anti-human IgM antibody (Jackson ImmunoResearch Laboratories, 109-035-129) was used at a 1:10,000 dilution (10µL/well) followed by the addition of SureBlue Reserve TMB substrate (KPL). Reactions were stopped by Stop Solution (KPL, Gaithersburg, MD) after five minutes and optical density (OD) was detected at 450 nm (Perkin Elmer, Victor). DH1017.IgM was used as a positive control and standard. Blank wells were used as a negative control. Antibody concentrations were interpolated from the standard curve, which was fit using to a 5-parameter fit sigmoidal curve (SoftMax Pro 6.3; Molecular Devices). Concentrations were interpolated from OD_450_ values within the linear range of the sigmoidal curve at dilution 90, or alternatively 30, for samples at 2-6 DPI, and at dilution 10 for samples at 8+ DPI. A passing quality control criterion of replicate OD_450_ values less than or equal to 25% variance was applied. Limit of detection for this assay was 0.08 µg/mL across assays, based on the linear range of the standard curve.

To measure concentration of IgM from human plasma, the same assay was applied with the following modifications. All original and IgM/mock depleted samples were diluted 1:30 relative to original plasma with 3-fold serial dilutions. Human IgM (Jackson ImmunoResearch Laboratories, 009-000-012) was used as positive control and standard. IgM concentration in sample was inferred based upon the 1:2430, or alternatively the 1:270 and 1:810 serum dilutions, which were all in the linear range of the standard curve.

### Biolayer interferometry

BLI assays were performed on the ForteBio Octet Red96 instrument at 25°C with a shake speed of 1,000 RPM. Goat Anti-Human Ig polyvalent (20 µg/mL; Thermo Fisher) was amine coupled to AR2G sensor tips as follows: sensor tips were activated with s-NHS/EDC (300s), incubated in Anti-Human Ig (600s), and quenched with ethanolamine (300s). DH1017.IgM, DH1017.IgG, and a negative control IgG (palivizumab) were diluted to 50µg/mL in PBS and captured on the biosensors (300s), followed by a PBS wash (60s). A baseline signal was recorded for 2 min in PBS (pH 7.4). Biosensors were then immersed into a two-fold dilution series of ZIKV E protein dimer in PBS (50-0.78µg/mL) to measure association (400s), followed by immersion into PBS to measure dissociation (600s). All binding profiles were corrected by double reference subtraction using the signal obtained in a PBS control (without E proteins) and the signal obtained from the Palivizumab control sensor. All affinities were calculated using the fast kinetic components of the heterogeneous ligand model fit. Kinetics and affinity data are the result of a single measurement.

### Size exclusion chromatography

ZIKV E dimer was screened for size on Superdex 200 Inc 10/300 GL column: 100uL of E dimer was loaded onto the column at 0.13mg/mL and was run at a flow rate of 0.5 mL/min in PBS.

### Virus propagation and purification for cryo electron microscopy

ZIKV strain H/PF/2013 was propagated in Vero-Furin cells (Mukherjee et al., 2016). Approximately 10^9^ Vero-Furin cells, grown in 10% FBS (Sigma, F0926), Dulbecco’s modified Eagle medium (DMEM) media (Thermofisher, 41300039) and 50 ug/mL blasticidin (Invivogen, anti-bl-1) at 37°C 5% CO_2_, were infected with a MOI of 0.1. Virus particles in 2% FBS/DMEM/1mM Pen/Strep (Thermofisher, 15140122) were incubated with cells for 2 hours at room temperature. After 2 hours, infected cells were incubated at 37°C, 5% CO_2_ for 36 hours. At 36 hours post infection (hpi), infectious media was collected and replaced with fresh media every 12 hours up to 84 hpi. Virus particles were purified from media collected at 60 and 72 hpi.

Briefly, cell debris were pelleted by centrifugation at 2,744 x g for 30 minutes. Infectious media was filtered with through a Steritop-GP 0.2 μm filter (Millipore, SCGPT05RE) and virus particles were precipitated overnight at 4°C with PEG 4000, 8% final concentration. PEG precipitated particles were concentrated at 8,891 x *g* for 50 minutes. Particles were concentrated through a 24% sucrose cushion by ultra-centrifugation at 126,144 x *g* for 2 hours at 4°C. Virus particles were isolated with a discontinuous potassium-tartrate/glycerol/NTE buffer (120mM NaCl, 20mM Tris pH 8.0, 1mM EDTA) gradient in 5% increments between 35% and 10% potassium-tartrate and ultra-centrifugation at 126,144 x *g* for 2 hours at 4C. The particles were extracted from the 20% fraction of the discontinuous gradient and buffer exchanged and concentrated in NTE buffer with a 100 kDa MWCO centrifugal filter.

### Single particle cryoelectron microscopy

Prior to freezing, purified virus and DH1017.Fab fragments were incubated on ice for 2 hours at a molar ratio of 0.2 mM Fab to 1 mM E protein. Samples were then plunge frozen in liquid ethane using a Cryoplunge 3 System (GATAN). Briefly, liquid nitrogen was used to liquefy ethane at −190°C. A 2.5µL volume of sample was spotted on lacey carbon grids (Ultrathin C on Lacey Carbon Support film, 400 mesh, Cu. Ted Pella, Inc. Product number 01824) and blotted with Whatman grade 1, 200mm circle filter paper (GE Healthcare Life Sciences, catalog number 1001-020) for 2.5 seconds. Blotting air pressure setting used was 100 psi.

Data were collected on a Titan Krios (Thermofisher) microscope equipped with a Gatan K3 detector using and Leginon software package (Suloway et al., 2005). A total of 1,929 cryo EM micrographs were collected with a nominal magnification of 64000x, 0.66 Å/pixel size, and an electron dose equivalent, of 35 e-/Å2. Motion correction and CTF calculations estimations were performed using MotionCorr2 (Zheng et al., 2017) and CTFFIND4 (Rohou and Grigorieff, 2015) respectively.

Automated particle selection picking (Sigworth, 2004) performed with cisTEM (Grant et al., 2018) selected 34,474 particles. A maximum-likelihood algorithm based 2D classification (Scheres et al., 2005; Sigworth, 1998) was performed with using cisTEM. From a subset of 2D classes, 4104 particles were selected for further processing.

Single particle reconstruction was performed according to the ‘gold standard’ method using jspr (Guo and Jiang, 2014). Briefly, the particles were divided equally into two randomly selected independent particle sets. Twenty ab-initio models were generated from a random set of 700 particles selected from the set of 4,104 particles. Two ab-initio models were selected, and one ab-initio model was assigned to one independent particle set and the other ab-initio model was assigned to the other independent data set. Each dataset was refined iteratively assuming icosahedral symmetry. Refinement resulted in two independent models that converged on the same structure. Following corrections for astigmatism, elliptical distortion, defocus, and the masking of the disordered nucleocapsid core, the final models of each independent data set were combined into a single final model. The resolution of the map was calculated at 0.143 from the FSC curve (Rosenthal and Henderson, 2003).

### Model fitting, refinement, and structure analysis

Models used in fitting viral structural proteins in the cryo EM density map are the E glycoprotein ectodomain of the ZIKV asymmetric unit (PDB 6CO18) (Sevvana et al., 2018) and a homology model of DH1017.Fab fragment. The Fab fragment model is composed of individual domains variable heavy, variable light, C, and lambda domains. The homology models of each Fab fragment domain were generated with I-TASSER (Roy et al., 2010; Yang et al., 2014; Zhang, 2008) and aligned with PYMOL (version 2.0, Schrodinger LLC) to an IgM cryoglobulin Fab crystal structure (PDB 2AGJ) (Ramsland et al., 2006) to form the complete DH1017.Fab homology model. The C_α_ backbones of all structures were fit in the cryo-EM map using Chimera (Pettersen et al., 2004) fit-to-map function. The position of the Fab domains was further refined with Coot (Emsley, P 2010). Residues of the E ectodomain C_α_ backbone within 6 Å of the Fab fragment C_α_ backbone were identified with PYMOL. These residues are defined as the Fab footprint and mapped to the surface of the virus particle with RIVEM (Xiao and Rossmann, 2007). The binding of a human IgM pentamer to the virus surface at the five-fold icosahedral axis was modeled. The solution structure of an IgM pentamer (PDB 2CRJ) (Perkins et al., 1991) and the homology model of DH1017.Fab from this paper were used to build the model. The Fc domains Cμ4, Cμ3, and Cμ2 of PDB 2CRJ were modeled to similar positions of Cμ4 (PDB 4JVW), Cμ3 (PDB 4BA8), and Cμ2 (4JVU) domains fit to the 25A tomogram of an IgM pentamer bound to the surface of an artificial liposome (Müller et al., 2013; Sharp et al., 2019). One Fab domain from each monomer of the pentamer structure PDB 2CRJ were fit to the density map after being aligned manually to the fitted DH1017.Fab structure.

### Assessing protection in ZIKV mouse model

Animal husbandry and experiments were performed under the approval of the University of North Carolina at Chapel Hill Institutional Animal Care and Use Committee. Five-week-old male *Ifnar1*^-/-^ mice were inoculated with a lethal dose of ZIKV strain H/PF/2013 (1 x 10^3^ FFU) subcutaneously in the footpad on day 0 (Lazear et al., 2016). On days −1 and +1, 100µg of antibody was delivered intravenously via the retro-orbital route. Serum was collected every two days for 11 days to monitor viremia by qRT-PCR and survival was monitored for 15 days. Animals that lost ≥30% of their starting weight or that exhibited severe disease signs were euthanized.

### Measuring viral burden in mice

RNA from the serum of ZIKV-infected mice was extracted with the Qiagen viral RNA minikit. ZIKV RNA was quantified by TaqMan one-step qRT-PCR using a CFX96 Touch real-time PCR detection system (Bio-Rad). Genome copies per mL of serum on a log_10_ scale was determined by comparison with a standard curve generated by using serial 10-fold dilutions of a ZIKV plasmid and previously reported primers: forward primer CCGCTGCCCAACACAAG, reverse primer CCACTAACGTTCTTTTGCAGACAT, and probe 5′-FAM (6-carboxyfluorescein)-AGCCTACCT-ZEN-TGACAAGCAATCAGACACTCAA-3′IABkFQ (Integrated DNA Technologies) (Carbaugh et al., 2019).

### Reactivity to autoantigens

ELISA was performed in 384-well plates (Corning) as previously described (Bonsignori et al., 2011). Briefly, plates were coated overnight at 4°C with 15μl purified proteins at optimized concentrations in 0.1M Sodium Bicarbonate: DNA (Worthington, LS002105) at 5μg/mL, Centromere B (Prospec, PRO-390) at 0.15μg/mL, Histone (Sigma, H9250) at 0.2μg/mL, Jo-1 (Immunovision, JO1-3000) at 0.05 units/well, RnP/Sm (Immunovision, SRC-3000) at 0.2 units/well, Scl-70 (Immunovision, SCL-3000) at 0.4 units/well, Sm (Immunovision, SMA-3000) at 0.1 units/well, SSA (Immunovision, SSA-3000) at 0.2 units/well, and SSB (Immunovision, SSB-3000) at 0.1 units/well. Subsequently, plates were blocked (50µL/well) with assay diluent (PBS containing 4% [w/v] whey protein, 15% normal goat serum, 0.5% Tween 20) for 2 hours at room temperature. For DNA, plates were pre-coated with Poly-L-Lysine (Sigma, P6282) at 2μg/mL in PBS overnight at 4°C, and the assay diluent was PBS containing 2% [w/v] bovine serum albumin and 0.05% Tween-20. Monoclonal antibodies were tested using 3-fold serial dilutions starting at 10μg/ml. 10μl of primary antibodies were added to each well and incubated for 1 hour at room temperature. The following positive control antibodies from Immunovision were all tested at a 1:25 starting dilution with 3-fold serial dilutions: Anti-Centromere B (HCT-0100), Anti-single stranded DNA (HSS-0100), Anti-Histone (HIS-0100), Anti-Jo 1 (HJO-0100), Anti-SRC (HRN-0100), Anti-Scl 70 (HSC-0100), Anti-Sm (HSM-0100), Anti-SSA (HSA-0100), and Anti-SSB (HSB-0100). The negative control was assay diluent alone. Plates were developed using 15μl/well of combination of horseradish peroxidase–conjugated antibodies in assay diluent comprising) goat anti-human IgG (Jackson ImmunoResearch Laboratories,109-035-098) at 1:10,000 dilution; 2) goat anti-human IgM (Jackson ImmunoResearch Laboratories,109-035-129) at 1:10,000 dilution; 3) goat anti-human H+L (Promega, W403B) at 1:3,000 dilution. After a 1 hour incubation, plates were developed with 20μl/well of SureBlue Reserve TMB substrate (KPL) and stopped by 20μl/well Stop Solution (KPL) after five minutes. Optical density (OD) was detected at 450 nm (Perkin Elmer, Victor).

### HEp-2 Cell Staining

Indirect immunofluorescence (Zeuss Scientific) binding of DH1017 mAbs to HEp-2 cells was performed as previously described (Bonsignori et al., 2014). IgG1 mAbs 2F5 and 17B were used as positive and negative controls, respectively. All mAbs were tested at 25ug/ml and 50ug/mL. Images were acquired for 8 seconds with a 40x objective.

