## Supplemental material for "Zika virus-specific IgM elicited during pregnancy exhibits ultrapotent neutralization"

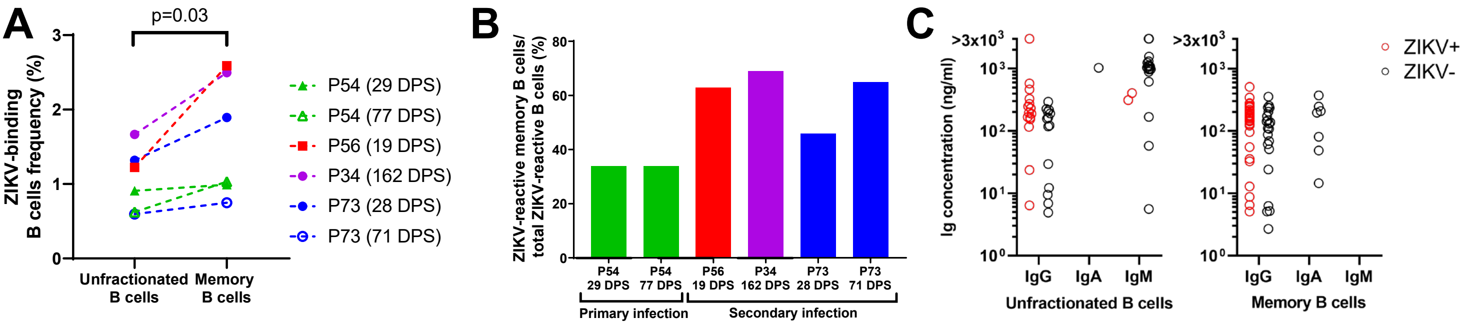


**Figure S1: Frequencies of ZIKV-reactive B cells.** Related to Table 1. **A.** Frequency of peripheral blood ZIKV-binding unfractionated and memory B cells. Days-post symptoms (DPS) of PBMC collection are indicated for each pregnant woman. **B.** Proportion of ZIKV-reactive memory B cells ($CD3^-/CD14^-/CD16^-/CD19^+/IgD^-/ZIKV^+$) over total B cells ($CD3^-/CD14^-/CD16^-/CD19^+/ZIKV^+$). **C** Immunoglobulin (Ig) concentration in supernatants of cultured unfractionated (left) and memory (right) B cells, clustered by isotype. Red circles indicate cultures for which ZIKV specificity was confirmed ($OD_{450}$ range= 0.44-2.5).

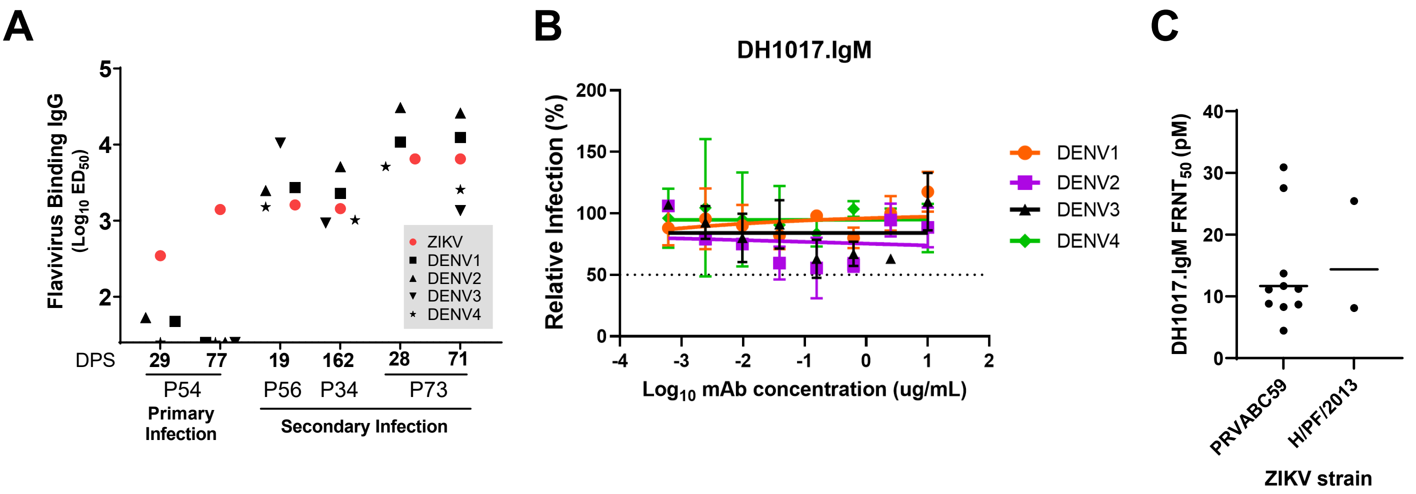


**Figure S2. Cross-reactivity with DENV 1-4 serotypes.** Related to Figure 2. **A.** ZIKV and DENV1-4 virions binding plasma IgG measured by ELISA and expressed as $Log_{10}$ $ED_{50}$ from all collections from which B cell responses were analyzed (n=6; days post symptoms, DPS). Maternal flavivirus serostatus as previously defined (Singh et al., 2019) is shown for each subject. Plasma samples were run in duplicate. **B.** DH1017.IgM neutralization by each of the four DENV serotypes. Bars indicate standard deviations of technical replicates run in duplicate. **C.** DH1017.IgM repeated neutralization assays against the ZIKV PRVABC59 (n=10) and H/PF/2013 (n=2) strains. Replicate experiments were independently performed by multiple operators and at different institutions for ZIKV H/PF/2013.

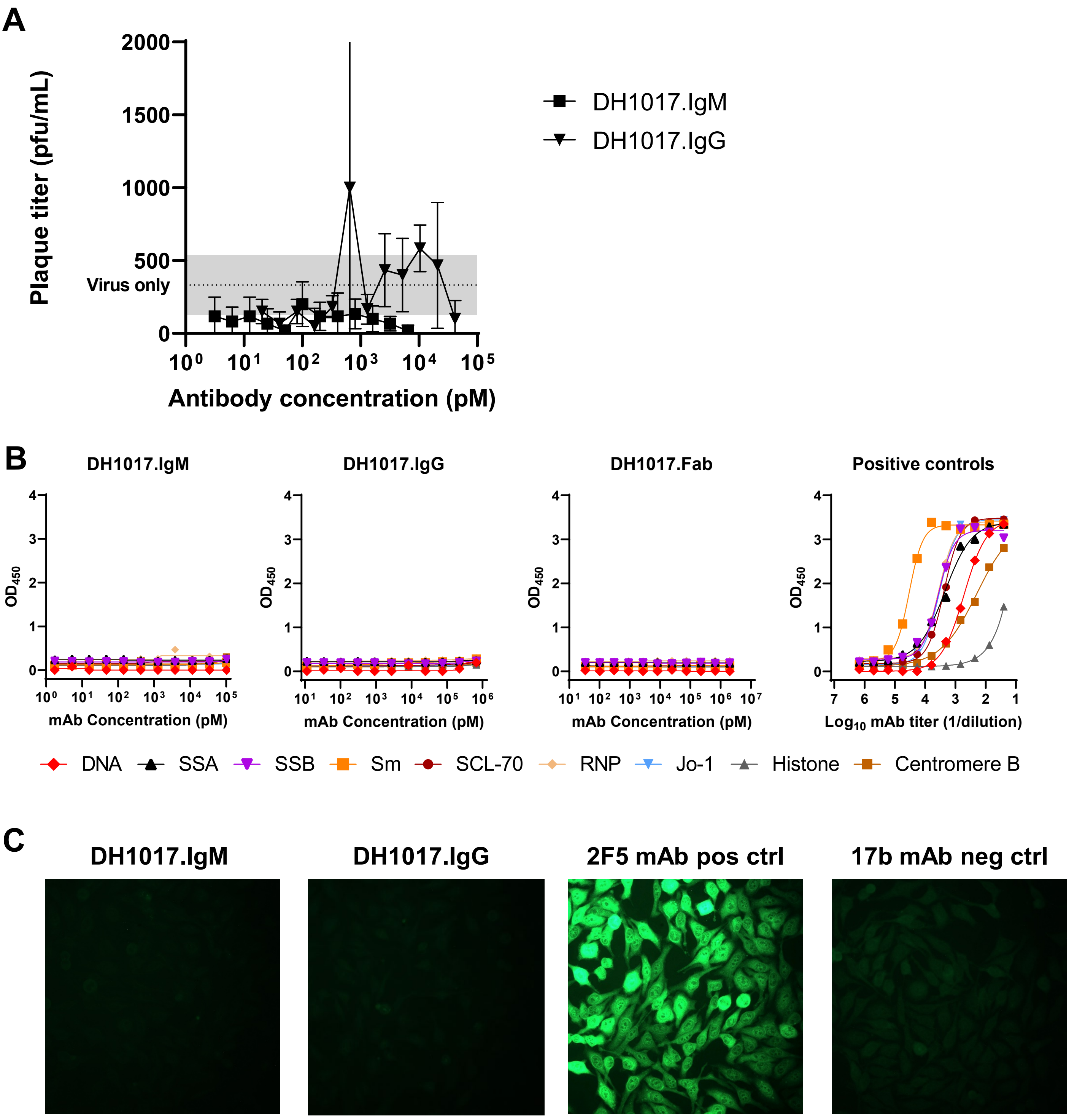


**Figure S3. Functional *in vitro* characterization of DH1017.IgM and DH1017.IgG.** Related to Figure 3. **A.** Antibody-dependent enhancement of DH1017.IgM and DH1017.IgG on primary monocytes measured in a plaque assay. Dotted line shows mean of six virus-only control replicates and the grey area indicates one standard deviation above and below this mean. Tests were run using 6 replicates. Bars show the standard deviation from 6 replicates. **B.** DH1017.IgM, DH1017.IgG and DH1017.Fab binding to a panel of autoantigens (Sjogren's syndrome antigens A and B [SSA and SSB, respectively], Smith antigen [Sm], ribonucleoprotein [RNP], centromere B [Cent B], histone, scleroderma 70 [Scl 70], Jo-1 proteins, and DNA) measured in ELISA. Respective positive controls are shown in the right panel. Error bars show standard deviation of duplicates. **C.** Hep2 cells were stained with DH1017.IgG and DH1017.IgM, 2F5 mAb (positive control), and 17B mAb (negative control) at a concentration of 50 μg/mL. Images shown were taken with a 40X objective with an 8s exposure and are representative of duplicate experiments.

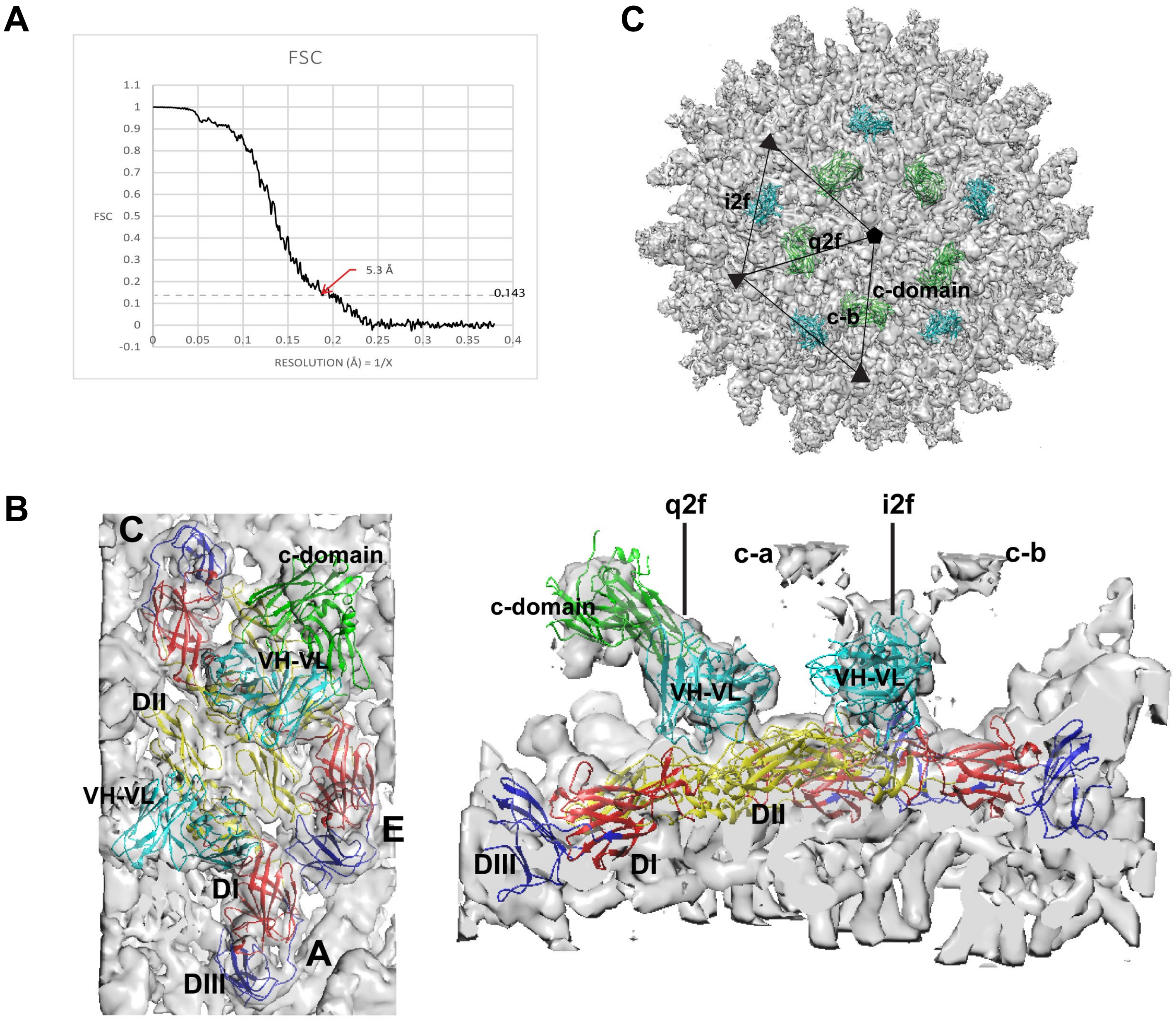


**Figure S4: DH1017.Fab interacts with ZIKV surface at multiple angles.** Related to Figure 4. **A.** FCS curve calculated from two half maps of the final model at 1.32 Å/pixel with the RNA core masked out. **B**. Section of electron density map fit with the asymmetric ectodomain, and a model of the DH1017 variable domain structure at the i2f axis and the Fab model at the q2f axis. DI, DII, and DIII are colored red, yellow, and blue, respectively. The DH1017 Fab fragment variable domain ($V_H$, $V_L$) is colored cyan, with the constant domain (c-domain) in green. The two different positions of constant domain density at the i2f axis are labeled c-a and c-b. The variable domain density at both axes is labeled $V_H$, $V_L$. The constant domain density at the q2f axis is labeled c-domain. The i2f and q2f axes are shown. **C.** Top-down view of i5f axis of density map with Fab DH1017 structures fit at the i2f axes, cyan structures, and q2f axes, green structures. Two neighboring asymmetric units are shown with open black triangles and the i5f and i3f axes are labeled as a pentagon and triangle, respectively.

**Table S1: Plasma IgM neutralization of ZIKV.** Related to Figure 1.

| ZIKV+ Subject ID | Maternal Serostatus | Days post symptoms | Sample type | IgM concentration (ug/mL) | Zika binding IgG (ED50) | ZIKV Neutralization (FRNT50) | Normalized ZIKV neutralization attributable to IgM (%) |
|---|---|---|---|---|---|---|---|
| 56 | 2° ZIKV | 14 | IgM dep | <LOD | 9433 | 3120 | 17 |
| | | | Mock dep | 62 | 11564 | 4594 | |
| 54 | 1° ZIKV | 36 | IgM dep | <LOD | 168.2 | 132.1 | 49 |
| | | | Mock dep | 113 | 198.1 | 303 | |
| 50 | 2° ZIKV | 36 | IgM dep | <LOD | 4304 | 412.4 | 19 |
| | | | Mock dep | NA | 2596 | 307.6 | |
| 50 | 2° ZIKV | 71 | IgM dep | <LOD | 3048 | 341.3 | 29 |
| | | | Mock dep | 100 | 2261 | 357.4 | |
| 54 | 1° ZIKV | 77 | IgM dep | <LOD | 359.9 | 859.4 | 37 |
| | | | Mock dep | 65 | 414.6 | 1582 | |
| 17 | 2° ZIKV | 166 | IgM dep | <LOD | 1365 | 400.5 | 0 |
| | | | Mock dep | 34 | 2923 | 690.9 | |
| 15 | 2° ZIKV | 178 | IgM dep | <LOD | 3977 | 3172 | 4 |
| | | | Mock dep | 26 | 4000 | 3335 | |
| 24 | 1° ZIKV | 186 | IgM dep | <LOD | 817 | 1479 | 0 |
| | | | Mock dep | 97 | 690.9 | 1257 | |
| 14 | 2° ZIKV | 188 | IgM dep | <LOD | 3402 | 1322 | 4 |
| | | | Mock dep | 43 | 3855 | 1561 | |
| 23 | 2° ZIKV | 209 | IgM dep | <LOD | 991.5 | 1327 | 28 |
| | | | Mock dep | 80 | 1869 | 3492 | |
| 19 | 2° ZIKV | 251 | IgM dep | <LOD | 1872 | 577.5 | 0 |
| | | | Mock dep | 31 | 2224 | 629.6 | |
| **Secondary ZIKV subject P73 with prolonged viremia in pregnancy** | | | | | | | |
| 73 | 2° ZIKV | 8 | IgM dep | <LOD | 14199 | 182.8 | 7 |
| | | | Mock dep | 39 | 17027 | 234.7 | |
| | | 14 | IgM dep | <LOD | 14544 | 290.3 | 78 |
| | | | Mock dep | 26 | 14569 | 1320 | |
| | | 28 | IgM dep | <LOD | 13457 | 626.2 | 30 |
| | | | Mock dep | 26 | 19377 | 1297 | |
| | | 42 | IgM dep | <LOD | 13153 | 232.7 | 41 |
| | | | Mock dep | 224 | 17824 | 538.3 | |
| | | 71 | IgM dep | NA | 7334 | 1340 | 52 |
| | | | Mock dep | NA | 9564 | 3623 | |
| | | 100 | IgM dep | <LOD | 5983 | 1315 | 3 |
| | | | Mock dep | 33 | 5658 | 1277 | |
| | | 112 | IgM dep | <LOD | 10132 | 1283 | 14 |
| | | | Mock dep | 18 | 12293 | 1804 | |
| | | 197 | IgM dep | <LOD | 9754 | 842 | 0 |
| | | | Mock dep | 21 | 9432 | 706.4 | |
| | | 289 | IgM dep | <LOD | 6020 | 392.5 | 0 |
| | | | Mock dep | 24 | 18170 | 708.3 | |
| | | 406 | IgM dep | <LOD | 6852 | 1939 | 0 |
| | | | Mock dep | 532 | 10584 | 1903 | |

**Table S2. Mutability analysis of nucleotide substitutions from germline in DH1017.IgM IgH and IgL V(D)J rearrangements.** Related to Table 1

$V_H$

| Region | Nucleotide mutation | Amino acid substitution | canonical hot/cold spot | s5f 5-mer model | Mutability rate |
|---|---|---|---|---|---|
| FR1 | a67g | T23A | - | gc**g**ct | 0.41 |
| CDR1 | g98a | G33D | g**g**tt | tg**g**tt | 1.30 |
| CDR1 | a101c | Y34S | t**a** | tt**a**ct | 1.74 |
| FR2 | a155c | Y52S | t**a** | gt**a**ca | 1.69 |
| FR3 | t214c | S72P | - | ta**t**ca | 0.46 |
| FR3 | g217a | V73I | a**g**ta | ca**g**ta | 4.19 |
| FR3 | c273t | D91 | - | ga**c**ac | 0.59 |

$V_L$

| Region | Nucleotide mutation | Amino acid substitution | canonical hot/cold spot | s5f 5-mer model | Mutability rate |
|---|---|---|---|---|---|
| CDR1 | a98g | Y33F | t**a** | tt**a**tg | 2.59 |
| FR2 | a116g | Q39R | - | gc**a**gc<br>gc***a***ac | 0.26<br>*1.16* |
| FR2 | g117a | Q39R | a**g**ct | ca**g**ct<br>c*g**g***ct | 2.21<br>*2.12* |
| FR2 | g130c | A44P | - | ca**g**cc | 0.21 |
| CDR2 | a155g | N52S | aa | ca**a**tg | 2.15 |
| CDR2 | a157g | N53D | - | gt**g**at | 1.34 |
| FR3 | c254g | A85G | - | gg**c**gg | 0.23 |
| CDR3 | c282g | S94R | - | agcag | 4.23 |
| CDR3 | c393t | A98V | tg**c**t | tgctg | 0.84 |

**Table S3. Epitope and paratope residues within 6Å between the DH1017.Fab and E ectodomain fitted structures to the cryo-EM density map of the Fab-bound Zika virion.** Related to Figure 4.

Fab footprint conserved residues at i2f and q2f symmetry axes

| Residue number(s) | Amino Acid | Domain/chain /axis |
|---|---|---|
| 120-126 | ACSKKMT | DII, A, C, i2f, q2f |
| 205,207,208 | T, N, N | DII, A, C, i2f, q2f |
| 230-232 | DTG | DII, A, C, i2f, q2f |
| 278-280 | DGA | DII, A, C, i2f, q2f |

Fab footprint unique residues at i2f and q2f symmetry axes

| Residue number(s) | Amino Acid | Domain/chain /axis |
|---|---|---|
| 61-66 | YEASIS | DII, A, i2f |
| 206,228,229 | M, G, A | DII, A, i2f |
| 277 | M | DI, A, i2f |
| 127,128 | G, K | DII, C, q2f |
| 233-235 | TPH | DII, C, q2f |
| 281 | K | DI, E, q2f |

Paratope footprint residues at i2f axis

| Residue number(s) | Amino Acid | Chain/region |
|---|---|---|
| 26-35 | GGSISSGDSY | VH, CDR1 |
| 54-59 | YYSGST | VH, CDR2 |
| 75-77 | TSK | VH, FR3 |
| 101-106 | VGDLRV | VH, CDR3 |

Paratope footprint residues at q2f axis

| Residue number(s) | Amino Acid | Chain/region |
|---|---|---|
| 26-28 | GGS | VH, CDR1 |
| 30-33 | SSGD | VH, CDR1 |
| 75-78 | TSKN | VH, FR3 |
| 101-106 | VGDLRV | VH, CDR3 |
| 57, 58 | G, I | VL, FR3 |
